# The effect of X-linked dosage compensation on complex trait variation

**DOI:** 10.1101/433870

**Authors:** Julia Sidorenko, Irfahan Kassam, Kathryn Kemper, Jian Zeng, Luke Lloyd-Jones, Grant W. Montgomery, Greg Gibson, Andres Metspalu, Tonu Esko, Jian Yang, Allan F. McRae, Peter M. Visscher

**Affiliations:** Institute for Molecular Bioscience, The University of Queensland, Brisbane, Australia; Estonian Genome Centre, Institute of Genomics, University of Tartu, Tartu, Estonia; School of Biology and Centre for Integrative Genomics, Georgia Institute of Technology, Atlanta, United States of America; Queensland Brain Institute, The University of Queensland, Brisbane, Australia

**Keywords:** Genome-wide association, dosage compensation, X chromosome, gene expression, complex traits, X inactivation

## Abstract

Quantitative genetics theory predicts that X-chromosome dosage compensation between sexes will have a detectable effect on the amount of genetic and therefore phenotypic trait variances at associated loci in males and females. Here, we systematically examine the role of dosage compensation in complex trait variation in humans in 20 complex traits in a sample of more than 450,000 individuals from the UK Biobank and in 1,600 gene expression traits from a sample of 2,000 individuals as well as across-tissue gene expression from the GTEx resource. We find, on average, twice as much genetic variation for complex traits due to X-linked loci in males compared to females, consistent with a negligible effect of predicted escape from X-inactivation on complex trait variation across traits and also detect biologically relevant X-linked heterogeneity between the sexes for a number of complex traits.

## Introduction

In eutherian mammals, including humans, females inherit two copies of the X chromosome and males only one. Ohno’s hypothesis posits that the dosage difference between the X chromosome and autosomes is resolved by doubling the expression of X-linked genes in both males and females, and to balance allele dosages differences in X-linked genes between the sexes, mechanisms have evolved to randomly inactivate one of the X chromosomes in females during embryogenesis, where female cells will express the maternal or paternal X chromosome approximately 50 percent of the time (Lyon, 1961; Ohno, 1967). X chromosome inactivation (XCI) is controlled by an approximately 1Mb region on the long arm of the X chromosome called the X inactivation centre. Initiation of the XCI process involves a step to ensure that at least two copies of the X inactivation centre are present in the female cell (Rastan and Robertson, 1985), and then the expression of the non-coding RNA X inactivation-specific transcript (*XIST*) from the X inactivation centre of the future inactive X chromosome (Brown *et al.*, 1991; Penny *et al.*, 1996; Panning, Dausman and Jaenisch, 1997). Rapid accumulation of *XIST* RNA is shown to start around the 8-cell human embryo development stage (van den Berg *et al.*, 2009) and most of female-to-male X-linked expression levels are equalized prior to embryo implantation (Petropoulos *et al.*, 2016; Moreira de Mello *et al.*, 2017). While exact dynamics of the human pre-embryonic XCI remain to be fully understood (Keniry and Blewitt, 2018), this process eventually resolves to the random transcriptional silencing of the one X chromosomes in female somatic cells. Random XCI remains maintained in mitotically derived cell lineages through a combination of epigenetic modifications including histone modifications and DNA methylation (Csankovszki, Nagy and Jaenisch, 2001; Lucchesi, Kelly and Panning, 2005) and leads to diverse patterns of mosaicism. However, approximately 15 to 23 percent of X-linked genes are shown to escape XCI (Carrel and Willard, 2005; Balaton and Brown, 2016; Tukiainen, A. Villani, *et al.*, 2017). Studies have previously used sex-bias in DNA methylation (Lister *et al.*, 2013; Cotton *et al.*, 2015; Schultz *et al.*, 2015) and gene expression (Johnston *et al.*, 2008; Zhang *et al.*, 2011) as an indication of XCI, where an inactivated X-linked gene in the non-pseudoautosomal region (non-PAR) of the X chromosome is expected to show no difference in expression between the sexes, while a non-PAR X-linked gene that escapes XCI is expected to have higher expression in females compared to males. Indeed, genes that show significant differences in expression between the sexes are enriched in escape genes, with the non-PAR region of the X chromosome enriched for genes with female-biased expression, and the PAR region enriched for genes with male-biased expression (Tukiainen, A.-C. Villani, *et al.*, 2017). The sex-bias in gene expression and its magnitude varies across tissues and even between the single cells, indicating variability in escape from XCI (Carrel and Willard, 1999; Tukiainen, A. Villani, *et al.*, 2017).

Sex is an important predictor for many quantitative traits, such as height, or the risk, incidence, prevalence, severity, and age-at-onset of disease (Ober, Loisel and Gilad, 2008). In addition to mean differences, males and females may also differ with respect to the trait variance (Lynch and Walsh, 1998). In this study, we focus on one aspect of Ohno’s hypothesis, where dosage compensation (DC) between the sexes is achieved by XCI. Theoretically, DC at loci affecting complex traits has a predictable effect on differences in genetic and therefore phenotypic trait variances in males and females and on the resemblance between male-male, male-female and female-female relatives (Bulmer, 1980; Lynch and Walsh, 1998; Kent, Dyer and Blangero, 2005). In particular, for X-linked complex trait loci, FDC is predicted to lead to twice as much variation in males compared to females and, conversely, escape from XCI is predicted to lead to twice the variance in females. Additionally, lack of DC can also contribute to mean differences in the trait of interest (Kent, Dyer and Blangero, 2005). Studies examining the relationship between X-linked SNPs and gene expression variation (Castagné *et al.*, 2011; Brumpton and Ferreira, 2016) and variation in complex traits (Zhang *et al.*, 2015) have noted that a larger proportion of SNPs are associated with these traits in males compared to females, indicating that these SNPs explain a larger proportion of variance in males compared to females. By comparing theoretical expectations from standard DC models to empirical data, we can systematically examine the effect of X-inactivation or escape from XCI on complex trait variation.

In this study, we leverage information on 20 complex phenotypes in the UK Biobank (N=208,419 males and N=247,186 females), 1,649 gene expression traits in whole-blood (N=1,084 males and N=1,046 females), and a mean of 808 gene expression traits across 22 tissue-types in GTEx (mean N=142 males and mean N=85 females) to compare the predicted effect of random X-inactivation in females to the empirical data. We perform a sex-stratified X-chromosome-wide association analysis (XWAS) for all traits to estimate male-female (M/F) ratio of the heritability attributable to the X chromosome in high-order UK Biobank traits and to compare M/F effect estimates of associated SNPs for both phenotypic and gene expression traits. Our results are consistent with expectations from full DC, and show a negligible effect of escape from XCI on complex trait variation.

## Results

### Evidence for dosage compensation in complex traits

We first performed a sex-stratified genome-wide association analysis for 20 quantitative traits in the UK Biobank (UKB) (for trait information see **Supplementary Table 1**), and estimated ratios of male to female SNP-heritabilities (*h*^*2*^_SNP_) on the X chromosome and the autosomes from summary statistics (Supplementary Material). Depending on the amount of DC on the X chromosome in females, this ratio is expected to take a value between 0.5 (no DC) and 2 (full DC). We refer to this as the DC ratio (DCR). For 19 out of 20 traits, the DCR estimates on the X chromosome (non-PAR) were significantly different from the expectation for no DC (DCR=0.5), and consistent with evidence for DC between sexes on the X chromosome and its detectable effect on phenotypic trait variation (**Figure 1A**, **black**). We validated our DCR summary statistics approach by calculating DCR from the estimates of *h*^*2*^_SNP_ in males and females derived from GCTA-GREML (Yang, Lee, *et al.*, 2011) on individual-level data from up to 100,000 unrelated individuals (**Supplementary Table 2**). From the GCTA-GREML analysis, we found the X-linked genetic variance of the complex traits to be low in general, but detectable in this large sample with the mean X-chromosome *h*^*2*^_SNP_ estimates of 0.62% (SD=0.34%) and 0.30% (SD=0.20%) across the 20 UK Biobank traits in males and females, respectively. These *h*^*2*^_SNP_ estimates were significant for all 20 traits in males and for 18 traits in females (the X-chromosome *h*^*2*^_SNP_ estimates for the skin and hair colour traits did not significantly differ from zero in the female-specific analysis) (**Supplementary Table 2**). For these 18 traits, we observe a strong overall correlation between DCR estimates obtained with the two methods (Pearson correlation, r=0.78) (**Supplementary Figure 1**).

**Figure 1:**
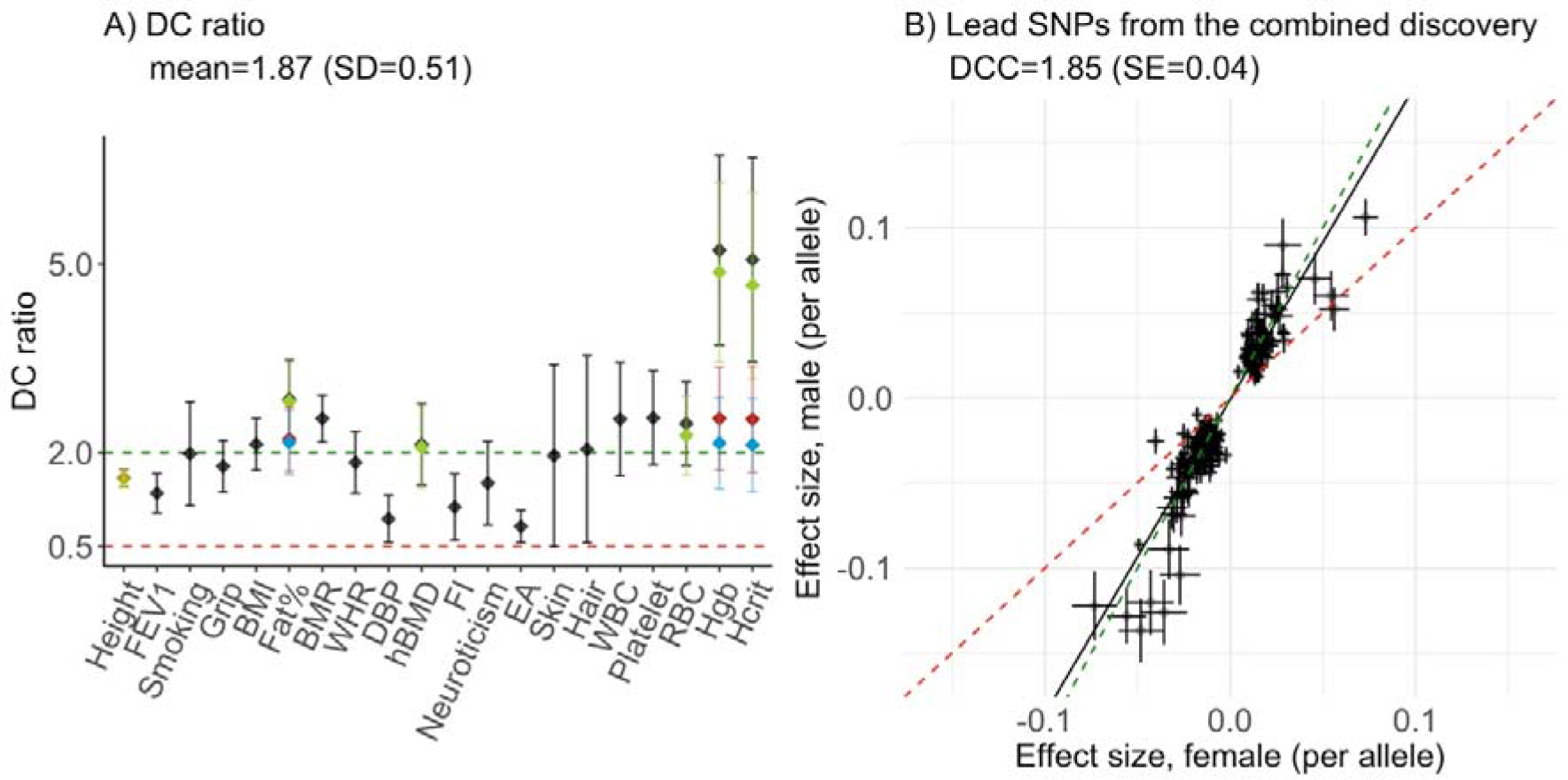
Estimates of DC ratio and dosage compensation coefficient for the UK Biobank traits. A) DC ratio with 95% confidence intervals (DC ratio +/− 1.96*SE) for 20 UKB traits as estimated using summary statistics from the association analyses. The estimates in black indicate the M/F ratio of the phenotypic variance explained by all SNPs on the X-chromosome (non-PAR). For height, Fat%, hBMD, RBC, Hgb and Hcrit the DC ratios are re-estimated excluding the SNPs in the regions of identified heterogeneity **(Supplementary Table 4)**and presented in colour (Excluding region 1=green; excluding region 2=yellow; excluding region 3 or 4=red; excluding region 1 and 3 or 4=blue). The mean DC ratio is estimated after accounting for heterogeneity. **B)**Male and female per-allele effect estimates (in standard deviation units) (+/− SE) are compared for the GWS SNPs identified in the combined discovery analysis (N=251). The SNPs located in the regions of heterogeneity for the six traits mentioned above are excluded. The green and red dashed lines indicate the expectations under full DC and escape from X-inactivation, respectively. The black line represents DCC. Height = standing height, FEV1 = forced expiratory volume in 1- second, Smoking = smoking status, Grip = hand grip strength (right), BMI = body mass index, Fat% = body fat percentage, BMR = basal metabolic rate, WHR = waist to hip ratio, DBP = diastolic blood pressure, hBMD = heel bone mineral density T-score, FI = fluid intelligence score, Neuroticism = neuroticism score, EA = educational attainment, Skin = skin colour, Hair = hair colour, WBC = white blood cell (leukocyte) count, Platelet = platelet count, RBC = red blood cell (erythrocyte) count, Hgb = haemoglobin concentration, Hcrit = Haematocrit percentage.

From the analysis based on summary statistics, the mean DCR for the X chromosome across 20 traits was 2.22 (SD=1.14), consistent with the expected value of 2 for full DC. In contrast, the estimates of the ratios of autosomal SNP-heritability varied from 0.66 to 1.17 with mean 0.95, in agreement with a limited difference in *h*^*2*^_SNP_ between the sexes in autosomal loci (**Supplementary Table 3**). We observed DCR on the X chromosome significantly different from expected values under both hypotheses (full and no DC) for nine traits (**Figure 1A**, **black**). While for standing height (height), forced expiratory volume in 1 second (FEV1), diastolic blood pressure (DBP), fluid intelligence (FI) and educational attainment (EA) the DCR estimates ranged between 0.5 and 2, indicating partial DC, values larger than 2 (body fat percentage (Fat%), basal metabolic rate (BMR), haemoglobin concentration (Hgb) and haematocrit percentage (Hcrit)) could not be explained under either of the DC models. We therefore sought an alternative explanation for these observations.

When estimating the DCR, we assumed that the genetic correlation (r_g_) between males and females is equal to one, and that any difference in the genetic variance is due to differences in dosage (i.e. number of active copies) of the X-linked genes. We estimated autosomal (r_gA_) and X-linked (r_gX_) genetic correlations in our sample using the GWAS summary statistics (see **Methods and Materials**). The evidence for autosomal genetic heterogeneity in complex trait is limited (Yang *et al.*, 2015; Rawlik, Canela-Xandri and Tenesa, 2016) and our estimates of r_gA_ between sexes are similar to published results (mean r_gA_=0.92, SD=0.06 across 20 traits, **Supplementary Table 3**). However, we found lower genetic correlation across the 20 traits on the X chromosome (r_gX_=0.80, SD=0.14) (**Supplementary Table 3**). The smallest r_gX_ estimates correspond to Hcrit (r_gX_=0.51, SE=0.05), Fat% (r_gX_=0.57, SE=0.05), red blood cell count (RBC) (r_gX_=0.64, SE=0.07) and Hgb (r_gX_=0.65, SE=0.04). These relatively low r_gX_ estimates may indicate local differences in genetic variance between males and females on the X chromosome that is independent of DC, which may explain the observed extreme DCR for these traits. We therefore explored biological heterogeneity as an explanation for these observations.

### Biological heterogeneity on the X chromosome

To investigate sex-specific genetic architectures on the X chromosome, we tested for heterogeneity in male and female SNP effects under the null hypothesis of no difference (**see Methods and Materials**). A total of 6 traits (Hcrit, Fat%, RBC, Hgb, height and heel bone mineral density T-score (hBMD)) showed evidence for heterogeneity, with four distinct heterogeneity signals. SNPs with significant differences in effect estimates between the sexes (P_Het_<5.0×10^−8^) were then LD-clumped to define four regions of heterogeneity, two of which overlap due to the complex LD structure in the centromere region (**Figure 2**, **Supplementary Table 4**).

**Figure 2:**
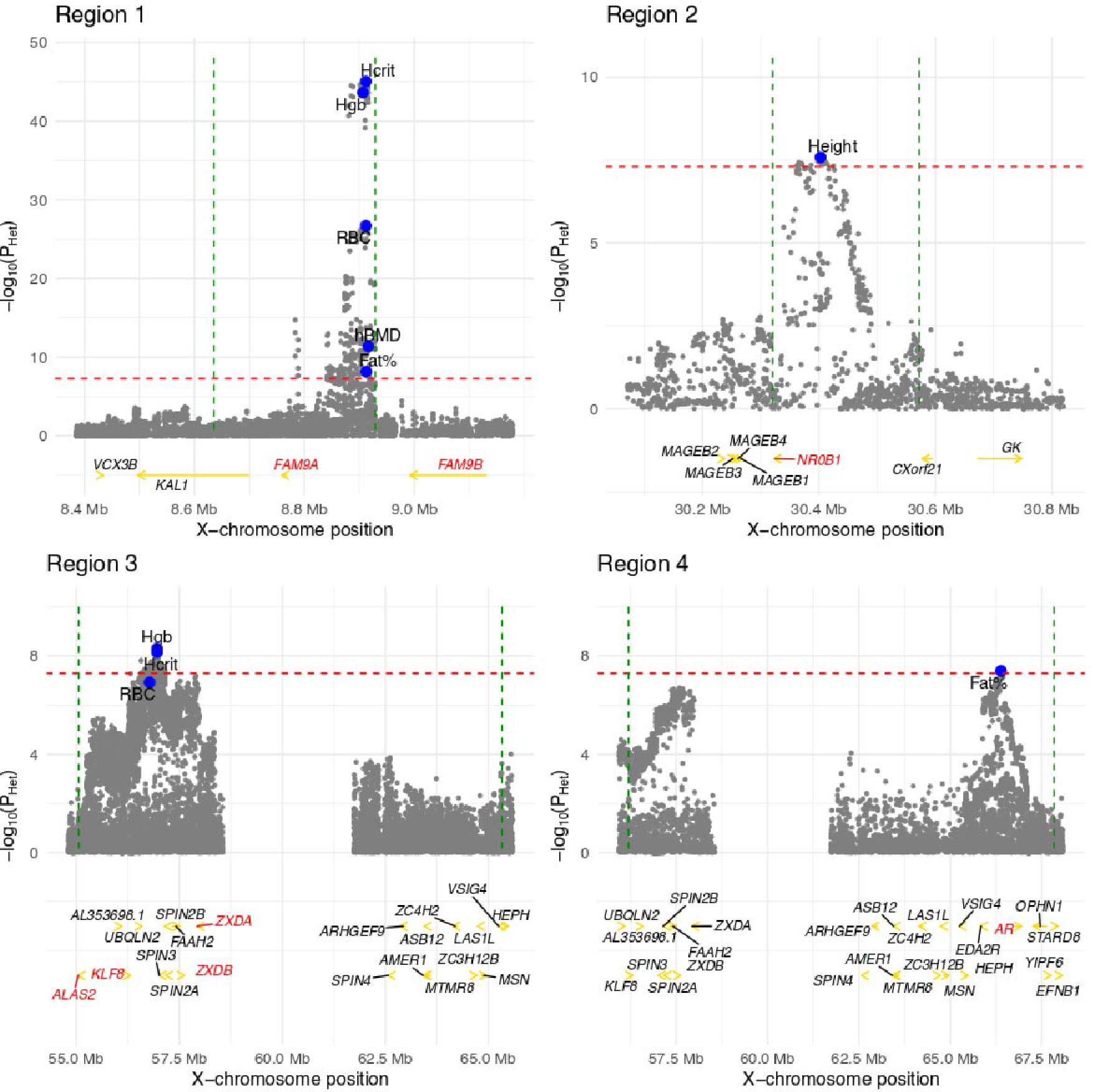
Four regions of heterogeneity (+/−250 kb) on the X chromosome. For each trait, regions of heterogeneity were identified as all SNPs within a region of LD R^2^ + 0.05 to the SNP with highest evidence of significant heterogeneity (**Supplementary Table 4**). In each region the P_Het_ values are plotted (grey dots) for all traits with significant heterogeneity in that region. The top SNPs for each trait are shown in blue. The genes discussed in the text are highlighted in red. In region 3, only the *ALAS2* gene and genes with X-chromosome position >56 Mb are shown for simplicity (the omitted 15 genes are: *ITIH6*, *MAGED2*, *TRO, PFKFB1*, *APEX2*, *PAGE2B*, *PAGE2*, *FAM104B*, *MTRNR2L10*, *PAGE5*, *PAGE3*, *MAGEH1*, *USP51*, *FOXR2*, *RRAGB*). The red dashed line represents the significance threshold (P_Het_ = 5.0×10^−8^). The green dashed lined represent the boundaries of the regions. P_Het_ = heterogeneity P-value, hBMD = heel bone mineral density T-score, Height = standing height, Hgb = haemoglobin concentration, Hcrit = Haematocrit percentage, RBC = red blood cell (erythrocyte) count, Fat% = body fat percentage.

Sex-related differences between males and females are most likely to arise due to naturally differing sex hormone levels. We therefore examined the evidence for hormonal regulation in these regions. We observed a highly significant trait association in males and lack of association in females in heterogeneity region 1 (Xp22.31) for 5 traits: Fat%, Hgb, Hcrit, RBC and hBMD (**Figure 2**). Notably, this region near the *FAM9A/FAM9B* genes, has been shown to be significantly associated male-specific traits such as testosterone levels (Ohlsson *et al.*, 2011), male pattern baldness (Pickrell *et al.*, 2016; Pirastu *et al.*, 2017) and age at voice drop (Pickrell *et al.*, 2016). Moreover, the *FAM9A/FAM9B* genes are shown to be expressed exclusively in testis in hybridization experiments (Martinez-Garay *et al.*, 2002). Indeed, in the GTEx data (see URLs), we found that *FAM9A* is highly expressed in testis only, with lower levels of expression of *FAM9B* in both uterus and testis, supporting the male-specific architecture for this locus and suggesting the androgenic pathway. Androgens play essential erythropoiesis promoting- (Shahani *et al.*, 2009), fat-reducing- (De Pergola, 2000) and anti-osteoporotic- (Clarke and Khosla, 2009) roles. Thus, we presume that a pleiotropic effect of the region 1 on erythropoiesis associated traits (Hgb, Hcrit and RBC), Fat% and hBMD may be mediated by androgen levels.

The *NROB1* gene in the region 2 (Xp21.2), which encodes the DAX1 protein, was a candidate gene for male-specific genetic control for height in this region (**Figure 2**). DAX1 is essential for regulation of hormone production and loss of DAX1 function leads to adrenal insufficiency and hypogonadotropic hypogonadism (Jadhav, Harris and Jameson, 2011). Moreover, Xp21.2 region in known as a dosage-sensitive sex reversal region, where its duplication or deletion is associated with male-female or female-male sex reversal (Bardoni *et al.*, 1994; Smyk *et al.*, 2007; Dangle *et al.*, 2017).

The top signal in region 4 was located in another well-known androgen-associated locus (Xq12) near the androgen receptor (*AR*) gene (**Figure 2**). The significant heterogeneity in this region between males and females for Fat% supports the male-specific fat-reducing effect of androgens. Notably, we observed the sex-specific heterogeneity in regions 1 and 4 for Fat% but not for BMI, suggesting that, although highly correlated, these traits differ in aetiology.

For hematopoietic traits (significant heterogeneity for Hgb and Hcrit, and nominal although not significant evidence for heterogeneity for RBC) the main heterogeneity signal was identified in Xp11.21 (region 3) (**Figure 2**). This region is shown to be associated with blood zinc concentrations (near *KLF8*, *ZXDA* and *ZXDB* encoding Zn-finger proteins (Evans *et al.*, 2013)) and male-pattern baldness (Pickrell *et al.*, 2016). Zinc has been shown to modulate serum testosterone levels in men (Prasad *et al.*, 1996) and is associated with haemoglobin concentrations in epidemiological studies (Houghton *et al.*, 2016). However, we find that the 5’ end of the region 3 is adjacent to the *ALAS2* gene, encoding a protein involved in heme synthesis and thus erythropoiesis (OMIM *301300). Mutations in this gene cause sideroblastic anaemia with X-linked recessive inheritance (OMIM #300751). Thus, the evidence for the androgen-dependent effect of this region on hematopoietic traits remains inconclusive.

Overall, at least three of the four regions of detected heterogeneity on the X chromosome show evidence of male-specific and/or androgen-related effects on the traits, and thus may not reflect an effect of DC, but rather biological differences between the sexes which are mediated by sex hormones. We therefore re-estimated DCR for Hcrit, Fat%, RBC, Hgb, height and hBMD after excluding these regions of heterogeneity (**Supplementary Table 5**, **Figure 1A**). While there was no significant change in DCR for height, we found a significant decrease in DCR and an increase in genetic correlation for the remaining five traits. After re-estimating DCR for the 6 traits our mean estimate of DCR across all 20 UK biobank traits changed from 2.22 (SD=1.14) to 1.87 (SD=0.51). These observations are consistent with the hypothesis that a disproportionate amount of male-specific genetic variance in these regions is at least partially hormonally influenced.

### Genetic effects of associated loci indicate limited escape from XCI in complex traits

In addition to testing for differences in overall X-linked variance between the sexes, we can estimate a dosage compensation parameter *d* such that *β*_*m*_ = *dβ*_*f*_ (see **Supplementary Methods and Material)**for genome-wide significant trait-associated SNPs. We did this by regressing the male-specific effect estimates onto the effects of the same markers estimated in female-specific analysis, weighted by the inverse of the variance of male-specific effect estimates. We define this regression slope as DC coefficient (DCC), which is expected to take on values between 1 (no DC or escape from XCI) and 2 (full DC).

We applied the conditional and joint association analysis (GCTA-COJO) (Yang *et al.*, 2012) to the summary statistics from the male-, female- and combined discovery analysis to select jointly significant trait-associated SNPs (hereafter, lead SNPs) for each of the 20 UKB traits. This identified 153 (male discovery) and 62 (female discovery) lead SNPs on the non-PAR X chromosome at a genome-wide significance level (GWS) (P<5.0×10^−8^) across the tested phenotypic traits (**Supplementary Table 6**-**8**). That is, more than twice the number of non-PAR lead SNPs was identified in males compared to females, indicating that a larger proportion of per-locus and therefore total genetic variance is explained in males compared to females. In contrast, in the PAR, we only identified two lead loci in males, while eight of them were detected in female discovery analysis (**Supplementary Table 6**-**9**). In the combined male-female discovery analysis 261 non-PAR and 16 PAR SNPs satisfy our GWS threshold in the COJO-analysis (**Supplementary Table 9)**. The increased number of lead SNPs in comparison to the sex-stratified analysis indicates concordance of effects from sex-specific analyses. The proportion of sex-specific genetic variance explained by the lead SNPs in the combined set is presented in **Supplementary Figure 2**.

We estimated DCC to be 2.13 (SE=0.08) and 1.46 (SE=0.08) for the male and female non-PAR discovery analyses, respectively, using the lead SNPs across the analysed complex traits (**Supplementary Figure 3)**. DCC for the markers identified in the combined analysis was 1.85 (SE=0.04) (**Figure 1B**). The observation from the combined analysis indicates only limited overall effect of escape from XCI on the variance or mean of the traits in our analysis. For the PAR, although the number of significant associations was small, the effects size estimates from sex-specific analyses were similar (**Supplementary Figure 4**), consistent with theoretical expectations.

The ratio of the M/F per-allele effect sizes for individual SNPs, which approximates the dosage compensation parameter, indicated the evidence for escape from XCI only for a few candidate variants. For instance, SNP rs113303918 in the intron of the *FHL1* gene is significantly associated with WHR in female and the combined analyses (P_*female*_=6.6×10^−12^ and P_*combined*_=9.8×10^−14^, respectively), while being only marginally significant in male-specific analysis P_*male*_=4.5×10^−5^) and the per-allele effect sizes on WHR are similar in both sexes (effect size ratio=0.93, SE=0.26). Similarly, the effect size ratio of SNP rs35318931 (P_*female*_=2.7×10^−17^, P_*male*_=6.7×10^−4^, P_*combined*_=2.8×10^−15^), a possible missense variant in the *SRPX* gene, is 0.63 (SE=0.20) consistent with escape from XCI for WHR. Assuming that these SNPs are the causal variants, the observed effect size estimates may indicate potential escape from XCI for *FHL1* and *SRPX*. Interestingly, for height (effect size ratio=2.12, SE=0.35; P_height, *combined*_=1.9×10^−37^) and BMR (effect size ratio=3.26, SE=1.21; P_BMR, *combined*_=6.6×10^−12^) the results for the SNP rs35318931 in the *SRPX* gene were indicative of DC. Consistent with these observations*, SRPX* is annotated with “Variable” XCI status in (Cotton *et al.*, 2013; Tukiainen, A.-C. Villani, *et al.*, 2017). For *FHL1*, although, annotated as “Inactive” in (Tukiainen, A.-C. Villani, *et al.*, 2017), findings from two earlier studies (Carrel and Willard, 2005; Cotton *et al.*, 2013), show that XCI is incomplete. Moreover, heterogeneous XCI of *FHL1* is detected in single cells and across tissues (Tukiainen, A.-C. Villani, *et al.*, 2017).

Previously, a locus near the *ITM2A* gene (SNP rs1751138, bp 78,657,806) was proposed as a potential XCI-escaping locus associated with height (Tukiainen *et al.*, 2014). In our sex-stratified and combined analyses from a sample size an order of magnitude larger, the lead marker for height was a nearby SNP rs1736534 located approximately 100 bp upstream of the previously reported rs1751138. The estimated M/F effect size ratio for the both variants was 1.75 (SE=0.11) (*β*_height, male_=−0.086, SE=0.004 and β_height, female_=−0.049, SE=0.002), providing evidence against extensive escape of *ITM2A* from XCI.

About one-third of the identified lead SNPs were physically located within X-linked gene regions. For these SNPs, we assigned the XCI status according to the reported XCI status of the corresponding genes (Tukiainen, A.-C. Villani, *et al.*, 2017) and compared the effect size ratios between “Escape/Variable” and “Inactive” genes. The results remained similar between two groups of genes (**Supplementary Figure 5**). A notable disadvantage of this approach is that the physical location of a SNP within a gene region does not necessarily indicate a causal variant for a complex trait. In contrast, an expression quantitative loci (eQTL) analysis avoids this, as there is no ambiguity between mapped SNPs and genes, and thus the annotation of XCI status.

### eQTL analysis indicates negligible escape from XCI in gene expression

We extended our DCC analysis to lower-order gene expression traits and performed a sex-stratified *cis*-eQTL analysis for 1,639 X-chromosome gene expression probes (28 of them in PAR) measured in whole blood. For each probe, we identified the top associated X-chromosome SNP with MAF>0.01 that satisfied the Bonferroni significance threshold of P<1.6×10^−10^ (i.e. 0.05/(1,639 × 190,245)) in the discovery sex (hereafter called eQTL), and extracted the same eQTL in the other sex and calculated DCC for M/F eQTL effect size estimates. We observed DCC of 1.95 (SE=0.04) for 51 eQTLs (48 unique SNPs) in the female discovery analysis, and DCC of 2.07 (SE=0.04) for 74 eQTLs (68 unique SNPs) in the male discovery analysis (**Supplementary Figure 6**), consistent with expectations from FDC and in agreement with our observations in high-order complex traits. We did not identify eQTLs for probes in PAR. Partitioning the non-PAR eQTLs based on reported XCI status of the corresponding genes (Tukiainen, A.-C. Villani, *et al.*, 2017) did not alter our results (**Figure 3**). In particular, for eQTLs annotated to escape XCI, DCC estimates were approximately two, consistent with FDC. Interestingly, for 6 eQTLs identified in the male discovery analysis and annotated to escape XCI (*USP9X*, *EIF2S3*, *CA5B*, *TRAPPC2*, *AP1S2*, and *OFD1*) we observed higher expression in females compared to males (P<3.1×10^−3^, i.e. 0.05/16), as expected for genes that escape from XCI, but found significant differences between the eQTL effect estimates of the top associated SNP on gene expression after correction for mean differences in expression between the sexes (genotype-by-sex interaction P<3.1×10^−3^), which is consistent with FDC. This suggests that sexual dimorphism in these genes may not be due to escape from XCI (**Supplementary Figure 7)**. Full details of the eQTLs in blood can be found in **Supplementary Tables 11 and 12**.

**Figure 3:**
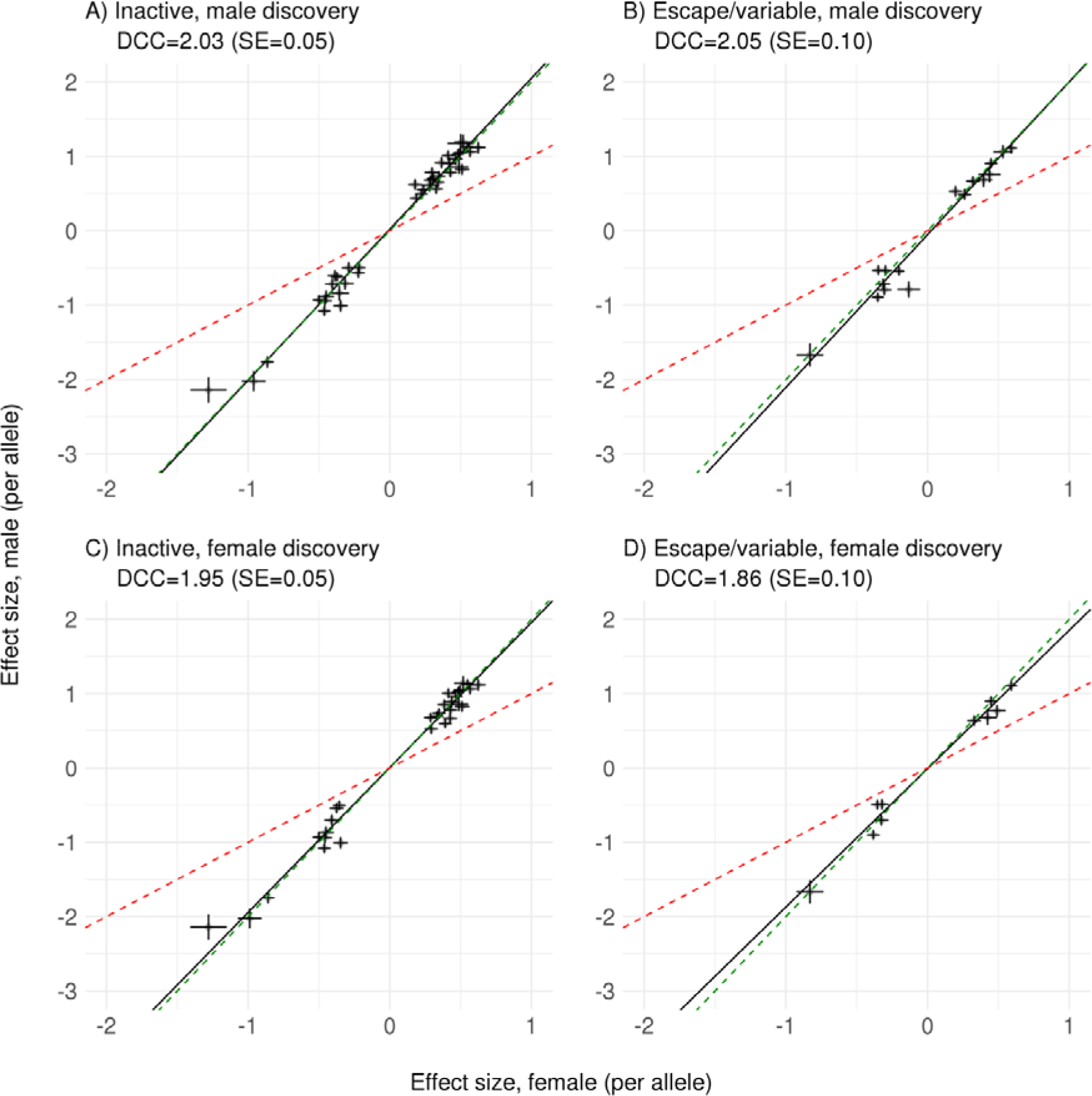
Dosage compensation coefficients for eQTLs from blood samples. A total of 62/74 and 45/51 eQTLs (P<1.6×10^−10^) in males and females, respectively, had either "Escape", "Variable", or "Inactive" status using annotations from (Tukiainen, A.-C. Villani, *et al.*, 2017). For 41 inactive eQTLs in the male discovery, DCC is 2.03 (SE=0.05), and for 16 escape or variable escape eQTLs, DCC is 2.05 (SE=0.10). For 30 inactive eQTLs in the female discovery, DCC is 1.95 (SE=0.05), and for 10 escape or variable escape eQTLs, DCC is 1.86 (SE=0.10). The red dashed line represents the expectation under escape from XCI. The green dashed line represents the expectation under FDC. The black line is the regression line.

We validated our results in 22 tissue samples from GTEx (v6p release) for which within tissue sample size was greater than N=50 in both males and females (**Supplementary Table 10**). We estimated DCC for at least three eQTLs (i.e. transcript-SNP pairs) that satisfied the within tissue Bonferroni significance threshold in the discovery sex in each of the 22 tissue-types. No eQTLs were identified for probes in PAR. A mean of 28 (SD=18) eQTLs were identified in the male discovery analysis across the 22 tissues. We observed a mean DCC of 1.94 (SD=0.16) across 22 tissues in the male discovery analysis, with the 95 percent confidence intervals for 20 tissues overlapping 2 (**Figure 4**). Heart (atrial appendage) tissue was an outlier, with DCC of 2.50 (SE=0.19). In contrast, a mean of 5 (SD=0.82) eQTLs were identified in females across the 7 tissues. A mean DCC of 1.59 (SD=0.13) across 7 tissues was observed in the female discovery analysis, with only the 95 percent confidence interval for thyroid tissue overlapping 2. We verified that the difference in estimated DCCs is not due to differences in sample size between males and females by down-sampling males so that the proportions match that of females within each of the 7 tissues and calculating mean DCC across 100 replicates (**Figure 4**). We did not observe enrichment for escape/variable eQTLs identified in the male or female discovery analyses by hypergeometric test (**Supplementary Table 11)**. These results were consistent when the top eQTLs were chosen among all tissues in the discovery sex and compared to the same eQTL from the same tissue in the other sex (**Supplementary Figure 8**). Finally, we compared our results to those from a sex-stratified autosomal *cis*-eQTL analysis in 36,267 autosomal gene expression probes in whole blood. A similar number of eQTLs with P<10^−10^ were identified in males and females (3,116 in the male discovery vs. 3,165 in the female discovery), indicating that an approximately equal proportion of autosomal genetic variance per locus is explained in each of the sexes. As expected, DCC in the male and female discovery was 1.00 (SE=2.3×10^−3^) and 0.94 (SE=2.3×10^−3^), respectively, indicating that the autosomal eQTL effect sizes are approximately equal in males and females (**Supplementary Figure 9**). Full details of the eQTLs across tissues can be found in **Supplementary Table 13**.

**Figure 4:**
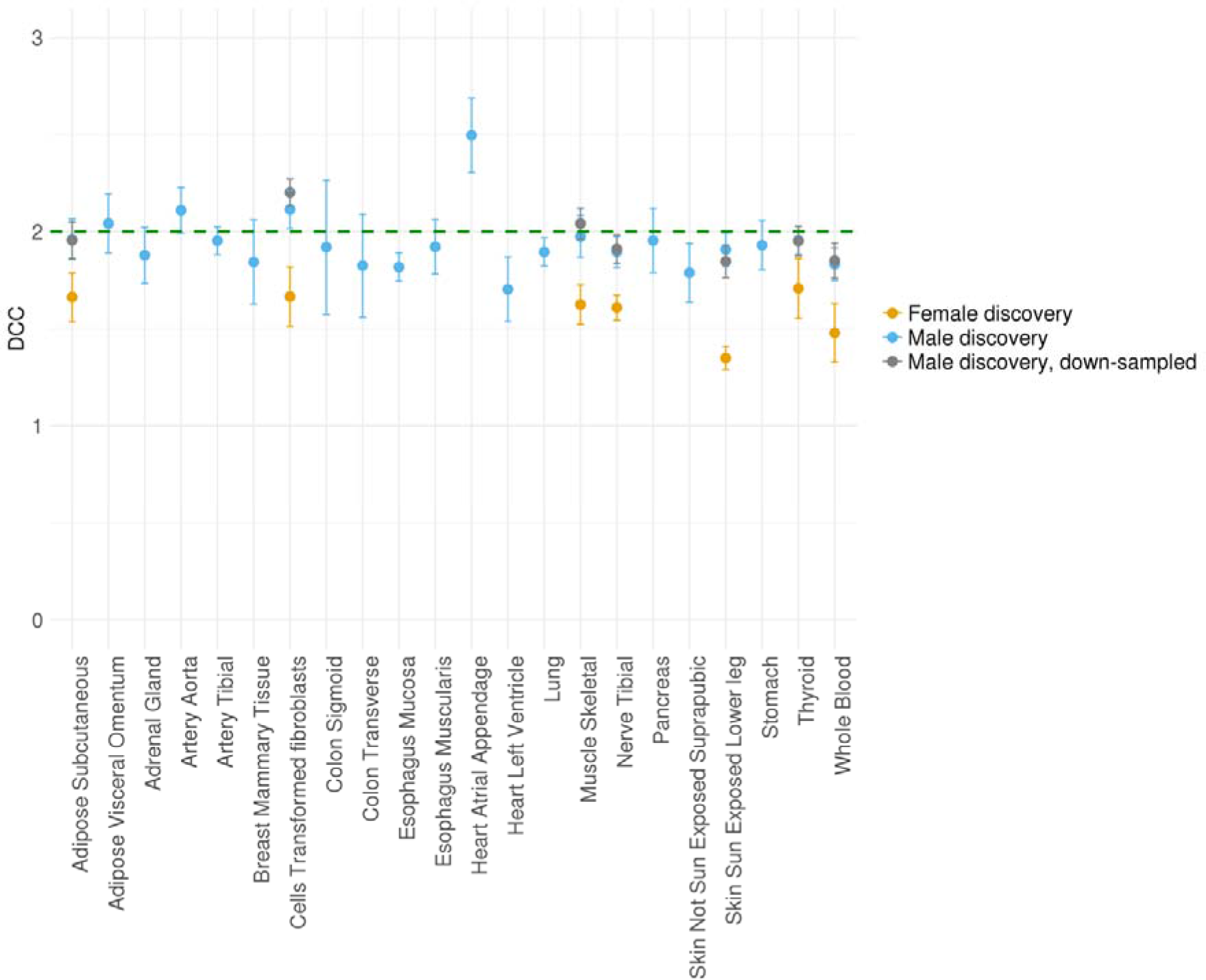
Dosage compensation coefficients for eQTLs across tissues. DCC is estimated for at least three eQTLs that satisfied the within tissue Bonferroni significance threshold in each of the 22 tissue-types. A mean of 27 (SD=17) eQTL are identified in the male discovery analysis giving a mean DCC of 1.93 (SD=0.20) across 22 tissues. A mean of 5 (SD=0.82) eQTLs are identified in the female discovery analysis giving mean DCC of 1.54 (SD=0.12) across 7 tissues. Males were down-sampled 100 times so that the proportions match that of females within each of the 7 tissues, and mean DCC is calculated across the 100 replicates. The bars represent the standard error.

### Summary-data based Mendelian randomisation

As noted above, there may be some ambiguity in mapping the associated variants to the genes based on its physical location, since the true causal variants may be masked by the local LD-structure or may exert the regulatory action on both near and distantly located genes (Smemo *et al.*, 2014; Zhu *et al.*, 2016). To investigate this we aimed to integrate the GWAS data from the complex trait analysis and the eQTL data from the whole blood analysis in the CAGE dataset to prioritize genes whose expression levels are associated with complex phenotypes because of pleiotropy, so that the XCI status would be assigned to the relevant “causal” gene. The combined summary data-based Mendelian randomisation (SMR) analysis (Zhu *et al.*, 2016) identified 18 genes (tagged by 20 probes) to be significantly (P_SMR_<3.0×10^−5^ (0.05/1,639) and P_HEIDI_>0.05) associated with 14 complex phenotypes (total of 37 associations) in the combined analysis (**Supplementary Table 14**). For males, associations between 13 genes (15 probes) and 11 traits satisfy our significance thresholds (total of 23 associations) (**Supplementary Table 15**), while for females we only identify 4 significant pleiotropic associations between 3 genes (3 probes) and 4 traits **(Supplementary Table 16**). The effects of the genetic variants on the trait, whose effects on the phenotype were identified to be potentially mediated by gene expression in sex-specific and combined analyses are shown in **Supplementary Figure 10.**The estimated DCC for these variants is similar to the results estimated with all jointly significant SNPs from COJO analysis (**Figure 1B, Supplementary Figure 3**).

Our SMR analysis linked many SNPs located in the intergenic regions to the expression of a number of genes, however, also a number of the SNPs physically located within a gene were determined to be associated with expression of another gene (e.g. a SNP in *TMEM255A* was an eQTL for *ZBTB33* whose expression is associated with traits skin and hair colour). This also included previous signals in escape genes being assigned to inactive genes (e.g. the SNPs physically located in the annotated escape gene *SMC1A* was associated with the expression of the inactive *HSD17B10* for BMI, BMR, Fat% and EA in the combined SMR analysis). Now the expression of only 2 genes (*MAGEE1* and *PRKX*) annotated with “Variable” or “Escape” (respectively) from XCI showed evidence for pleiotropic association with a phenotypic trait (hand grip strength (Grip) and white blood cells (WBC), respectively) due to a shared genetic determinant (*MAGEE1*: P_SMR,combined_=2.1×10^−6^, *PRKX:* P_SMR,combined_=8.7×10^−6^, **Supplementary Figure 10, Supplementary Table 14**). The estimated effect size ratio (2.84, SE=0.85) for the variant rs757314 (mediated by *MAGEE1* expression levels) on hand grip strength was not consistent with the escape from X-inactivation (the expected ratio for an escape gene is 1). For the rs6641619 (associated with *PRKX* expression and WBC), we estimate the effect size ratio of 1.33 (SE=0.44), which is indicative of partial escape from X-inactivation.

Variants near *ITM2A* were shown to be associated with height (Tukiainen *et al.*, 2014) and with height, BMR, Grip, WHR and FEV1 in the current study. In the combined SMR analysis we also observed evidence for pleiotropic association (P_SMR_<3.0×10^−5^) of the *ITM2A* (tagged by ILMN_2076600) expression with 7 traits: height, BMR, Grip, WHR, FEV1, DBP and RBC (genetic instrument rs10126553). However, only for the DBP and RBC, this association passes the test for heterogeneity (HEIDI), aimed to distinguish pleiotropy/causality from linkage. For the remaining traits, P_HEIDI_ varied from 6.5×10^−3^ for WHR to 8.0×10^−16^ for height, suggesting heterogeneity in gene expression effect on the trait estimated at different eSNPs that are in LD with the top-associated eSNP. That is, we cannot reject the null hypothesis that the gene-trait association is due to a single genetic variant. SMR analysis in *trans* regions on the X chromosome identified additional association between the expression of the *ITM2A* gene and height and BMR, which was mediated by a *trans*-eQTL located 2.2Mb upstream *ITM2A* (rs112933714). The mean M/F effect size ratio for the genetic instrument rs10126553 (P_eQTL,combined_=1.5×10^−76^) across these 7 traits (not filtered on P_HEIDI_ value) was 1.83 (SD=0.25) **(Supplementary Table 17)**, and 2.30 (SD=0.65) for the *trans* acting variant rs112933714 across two traits with significant *trans*-eQTLs **(Supplementary Table 18)**, in agreement with reported “Inactive” status of the *ITM2A* gene.

## Discussion

The theoretically predicted effect of random X-inactivation in female cells is two-fold reduced amount of additive genetic variance in females compared to males, whereas escape from XCI would increase genetic variance in females and contribute to sexual dimorphism. Having analysed phenotypes with varying degree of polygenicity, we found only limited effect of escape from X-inactivation on complex trait variation both in moderately (gene expression) and highly polygenic traits (phenotypic traits in the UKB). The two strategies that we use to estimate DC are the overall ratio of M/F X-linked heritabilities (i.e. the dosage compensation ratio) and the comparison of the individual effects of the trait-associated variants (i.e. the effect size ratio and dosage compensation coefficient). These are parameterisations of the same effect, the former based upon the variance contributed by all X-linked trait loci and the latter based upon per-allele effect sizes of trait-associated loci. Previous studies demonstrate that ~1% of phenotypic variance of the phenotypic traits, such as height and BMI, is attributable to the X chromosome (Yang, Manolio, *et al.*, 2011; Tukiainen *et al.*, 2014). However, the attempts to disentangle the relationships of additive genetic variance between the sexes in high-order traits were limited in power due to moderate sample sizes and/or computational challenges (Yang, Manolio, *et al.*, 2011; Tukiainen *et al.*, 2014). Here, a large the sample of > 205,000 males and >245,000 females allowed us to identify a statistically significant contribution of the X chromosome to the total trait heritability for 18 of the 20 studied complex traits in both sexes and for all traits in male-specific analysis, so we could make further inferences about DCR in complex traits. While we observed good overall evidence for DC across the phenotypic traits, a number of outliers were present in our analysis. First, we observed unexpectedly high ratios of male to female genetic variance for some of the traits. The male-specific genetic control for some genome regions appear to be sex-hormone dependent and thus are not informative on DC. Additionally, while the region comprising a testosterone-associated locus (near *FAM9A/FAM9B* genes) had the strongest evidence of heterogeneity, its removal had modest effect on DCR, while the exclusion of the genomic region near the centromere had the strongest effect. In addition to possible androgen-specific influence of this region, the tight LD structure could contribute disproportionately to sex-specific genetic variance. Second, we observe DCR supporting possible escape from XCI rather than full DC in brain related traits, such as educational attainment and fluid intelligence, and also diastolic blood pressure. Consistently, brain tissues have the highest X chromosome to autosome expression ratio, followed by heart (Nguyen and Disteche, 2005; Xiong *et al.*, 2010), in agreement with an enhanced X-chromosome role in cognitive functions. Thus, the effect of DC may be tissue-specific.

We also found consistent evidence for DC when examining individual trait-associated markers. Interestingly, our results for height associated loci near *ITM2A*, a gene known to be involved in cartilage development, differ from reported evidence for lack of DC (Tukiainen *et al.*, 2014) and only a few loci associated with WHR were candidates to be putative “escapees”. It should be noted, however, that for WHR genetic correlation on both autosomes and the X chromosome is markedly low, which may reflect the sex-specific genetic control for this trait.

In contrast to the complex phenotypic traits, gene expression has a notably different genetic architecture with as much as 65% of the expression variance for a gene explained by a single SNP alone, thus potentially violating the (polygenic) modelling assumptions for a DCR analysis, and thus was not included as part of this study. However, we were able to leverage information from eQTLs to show that DCC estimates in gene expression are consistent with expectations from FDC and in agreement with our observations in high-order complex traits and previous eQTL studies (Castagné *et al.*, 2011; Brumpton and Ferreira, 2016). These results were broadly consistent across multiple tissue-types, where a larger number of eQTLs were identified in males compared to females and, in the male discovery analysis, DCC is approximately 2. Across both the high-order and gene expression traits, we observed DCC estimates larger than 2 in the male discovery analysis and smaller than 2 in the female discovery analyses. This may be attributed to a combination of partial escape from XCI and “winner's curse” of the XWAS analysis. For example, any loci that partially escapes XCI in females would be preferentially selected in the female discovery analysis due to increased statistical power of detection, and thus bias the DCC estimates towards 1. Further, DCC estimates may be influence by “winner’s curse”, where the per-allele effect estimates in the discovery sex is biased upwards compared to the corresponding estimates in the other sex.

While identification of new associations between X-linked SNPs and complex traits was not the aim of our study, our results show these are readily found and that they cumulatively contribute to trait variation. For example, we find pleiotropic association between expression levels of the *HSD17B10* gene, which encodes a mitochondrial enzyme involved in oxidation of neuroactive steroids, fatty acids as well as sex hormones and its deficiency is implicated in neurodegenerative disorders (S. Y. Yang *et al.*, 2014) with obesity-related traits (Fat% and BMI) and educational attainment. Consistently, similar putative causal relationships were recently identified for the autosomal gene *HSD17B12*, where its increased expression of this gene was associated with decreased BMI across 22 tissues (Yengo *et al.*, 2018). Therefore, comprehensive surveys of sex-stratified X chromosome wide association studies for disease and other traits are likely to be rewarding, and may provide insight into new biology and sexual dimorphism. Moreover, since our method for estimating the amount of DC only requires summary statistics from association analyses, the availability of sex-stratified results from XWAS studies can further be informative on the effect and dosage of X-linked variation across a range of complex traits.

## Acknowledgements

This research was supported by the Australian Research Council (DP160101343, DP160102400 and DP160101056), the Australian National Health and Medical Research Council (1107258, 1078037, 1078399, 1113400, 1078901, 1083656, 1107258 and 1113400) and the US National Institutes of Health (R01 MH100141 and R21 ES025052). JY is supported by the Sylvia & Charles Viertel Charitable Foundation. The content is solely the responsibility of the authors and does not necessarily represent the official views of the grant funding bodies. This study makes use of data from GTEx Consortium data from dbGaP (accession phs000424.v6.p1). Complex trait analysis has been conducted using the UK Biobank Resource under project 12514. We thank Prof. Naomi R. Wray for her helpful comments and suggestions for the manuscript.

## URLs

GTEx, see https://www.gtexportal.org/home/.GCTA, see http://cnsgenomics.com/software/gcta/. SMR, see http://cnsgenomics.com/software/smr/. PLINK, see https://www.cog-genomics.org/plink2/. BOLT-LMM, see https://data.broadinstitute.org/alkesgroup/BOLT-LMM/. UK Biobank, see http://www.ukbiobank.ac.uk/. BioMart, see http://grch37.ensembl.org/biomart/martview/.

## Author Contributions

PMV and AFM conceived and designed the project. PMV, GWM, GG, AM, and TE conceived and designed the gene expression experiments and provided gene expression data. JS and IK performed the statistical analyses. JS, IK, KK, JZ, and LLJ performed the data quality control. JY contributed to the critical discussion and interpretation of the results. JS, IK, AFM and PMV wrote the manuscript. All authors read and approved the final manuscript.

## Declaration of Interests

We declare that we have no competing interests.

---

**Supplementary Figure 1.**
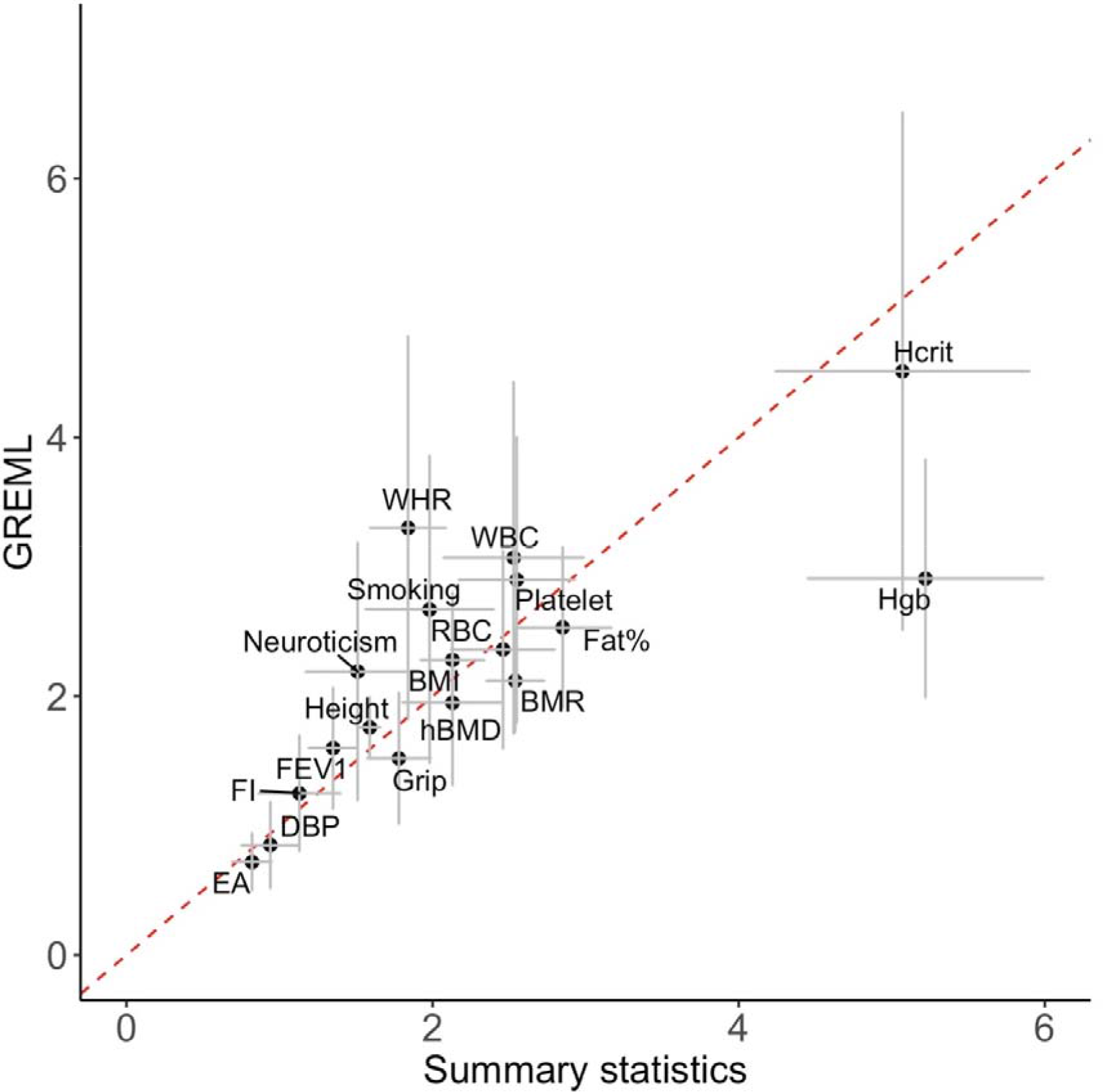
DC ratio estimates from summary statistics and REML for 18 traits with significant REML heritability estimates on the X chromosome in both sexes. The red dotted line indicates the expected correlation of 1. (Height = standing height, FEV1 = forced expiratory volume in 1-second, Smoking = smoking status, Grip = hand grip strength (right), BMI = body mass index, Fat% = body fat percentage, BMR = basal metabolic rate, WHR = waist to hip ratio, DBP = diastolic blood pressure, hBMD = heel bone mineral density T-score, FI = fluid intelligence score, Neuroticism = neuroticism score, EA = educational attainment, Skin = skin colour, Hair = hair colour, WBC = white blood cell (leukocyte) count, Platelet = platelet count, RBC = red blood cell (erythrocyte) count, Hgb = haemoglobin concentration, Hcrit = Haematocrit percentage)

**Supplementary Figure 2.**
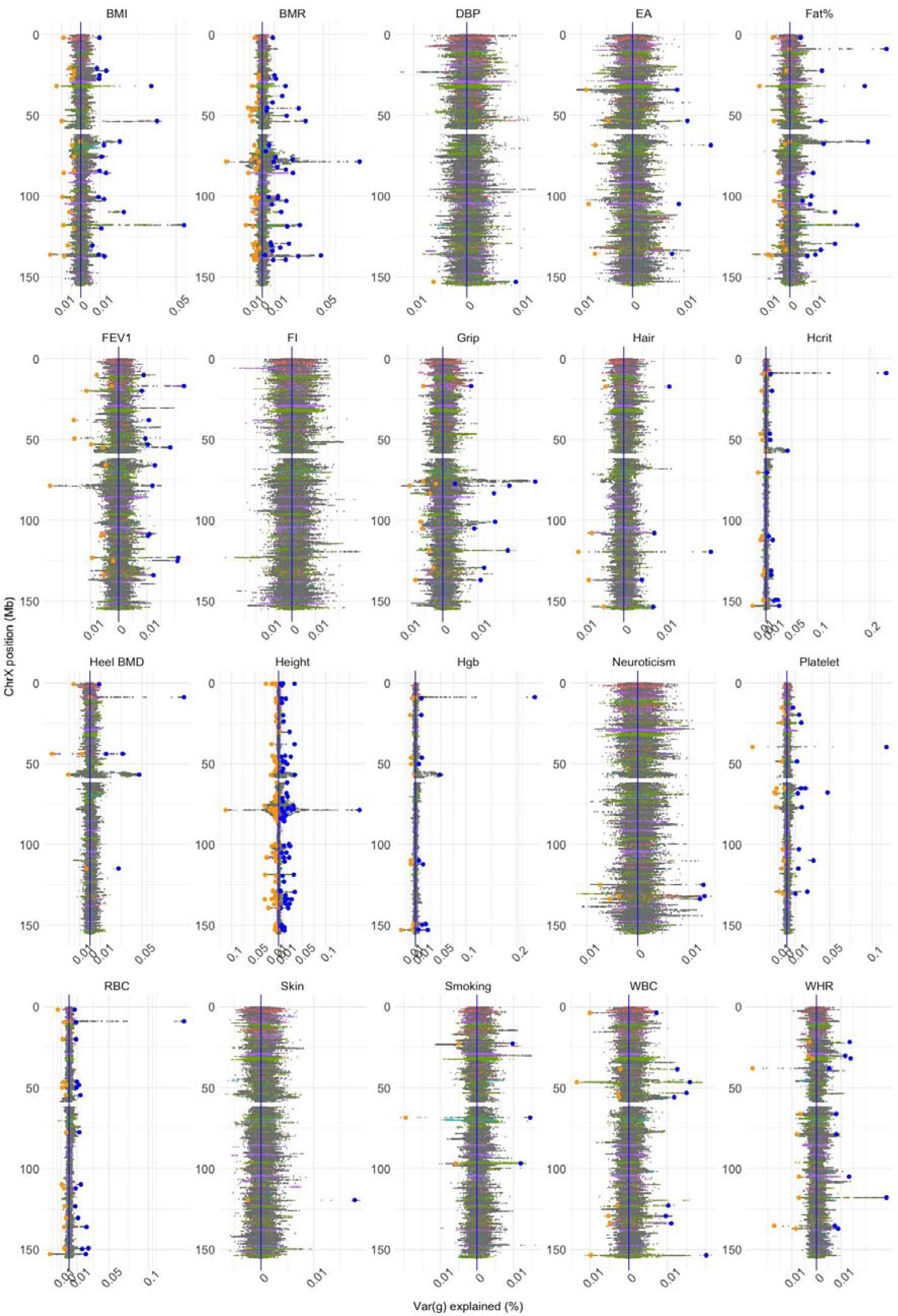
Sex-specific variance explained on the X chromosome. Genetic variance contributed by the SNP is each sex was calculated as 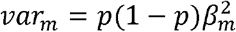 and 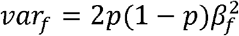 and 2*p*(1 − *p*)*β*^2^ for the SNPs in the PAR region. The per-allele effect estimates are from sex-stratified XWAS analysis. Sex-specific variance of the lead SNPs selected in the combined COJO-GCTA analysis are highlighted by larger circles (Blue colour represents males and orange - females). The base pair positions with the reported inactivation status (Tukiainen, A.-C. Villani, *et al.*, 2017) are highlighted in colour as follows: “Escape” - red, “Variable” -purple, “Inactive” -green, “Unknown”- light blue, “Non-available” (NA) - grey. (Height = standing height, FEV1 = forced expiratory volume in 1-second, Smoking = smoking status, Grip = hand grip strength (right), BMI = body mass index, Fat% = body fat percentage, BMR = basal metabolic rate, WHR = waist to hip ratio, DBP = diastolic blood pressure, heel BMD = heel bone mineral density T-score, FI = fluid intelligence score, Neuroticism = neuroticism score, EA = educational attainment, Skin = skin colour, Hair = hair colour, WBC = white blood cell (leukocyte) count, Platelet = platelet count, RBC = red blood cell (erythrocyte) count, Hgb = haemoglobin concentration, Hcrit = Haematocrit percentage)

**Supplementary Figure 3.**
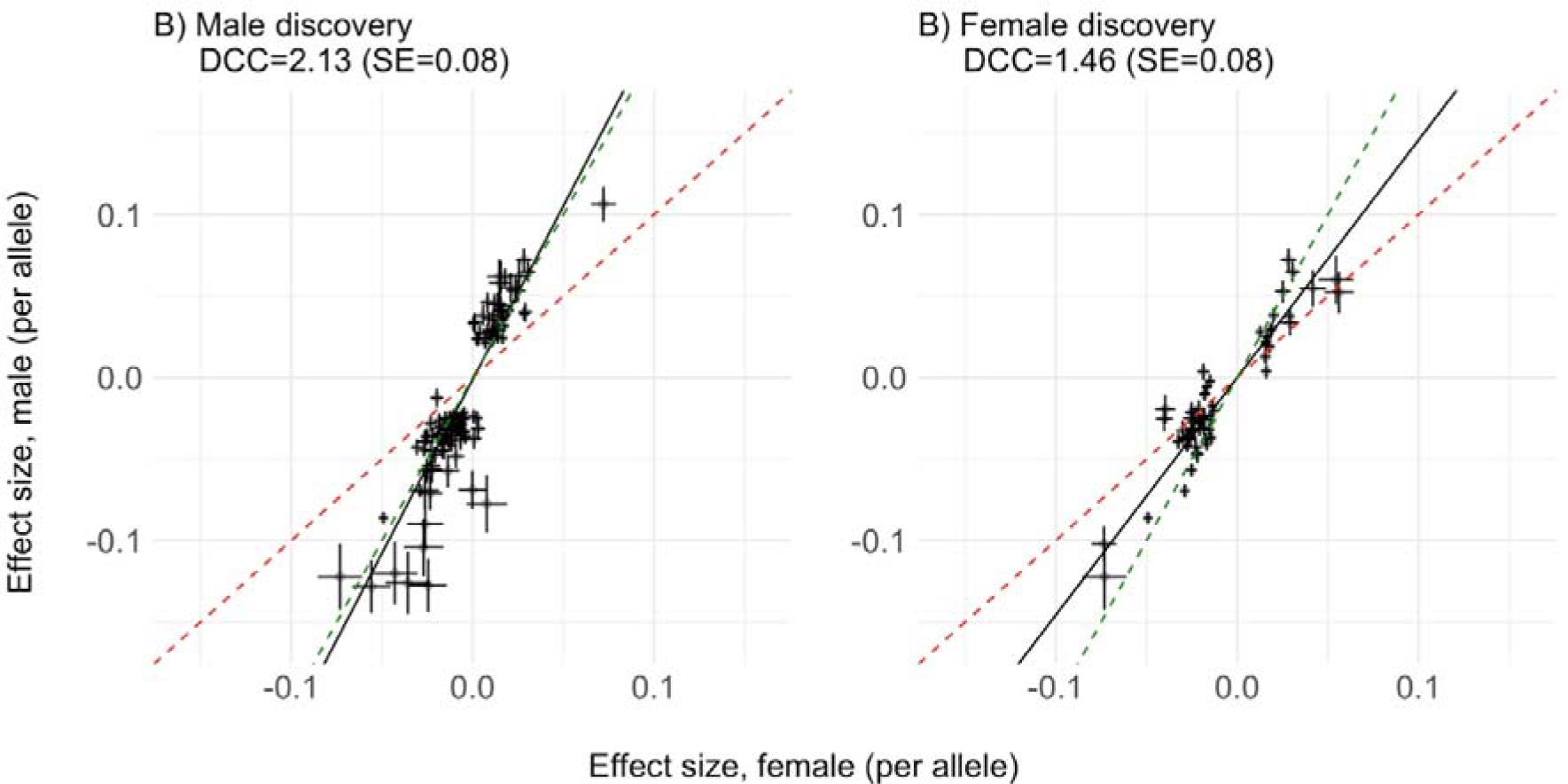
Comparison of the male- and female-specific per-allele effect estimates (+/− SE) for the lead SNPs (non-PAR) identified in the **B)**male discovery set (N=143) or **C)**female discovery set (N=62). The SNPs located in the regions of heterogeneity are excluded. The green and red dashed lines indicate the expectations under full DC and escape from X-inactivation, respectively. The black line represents DCC.

**Supplementary Figure 4.**
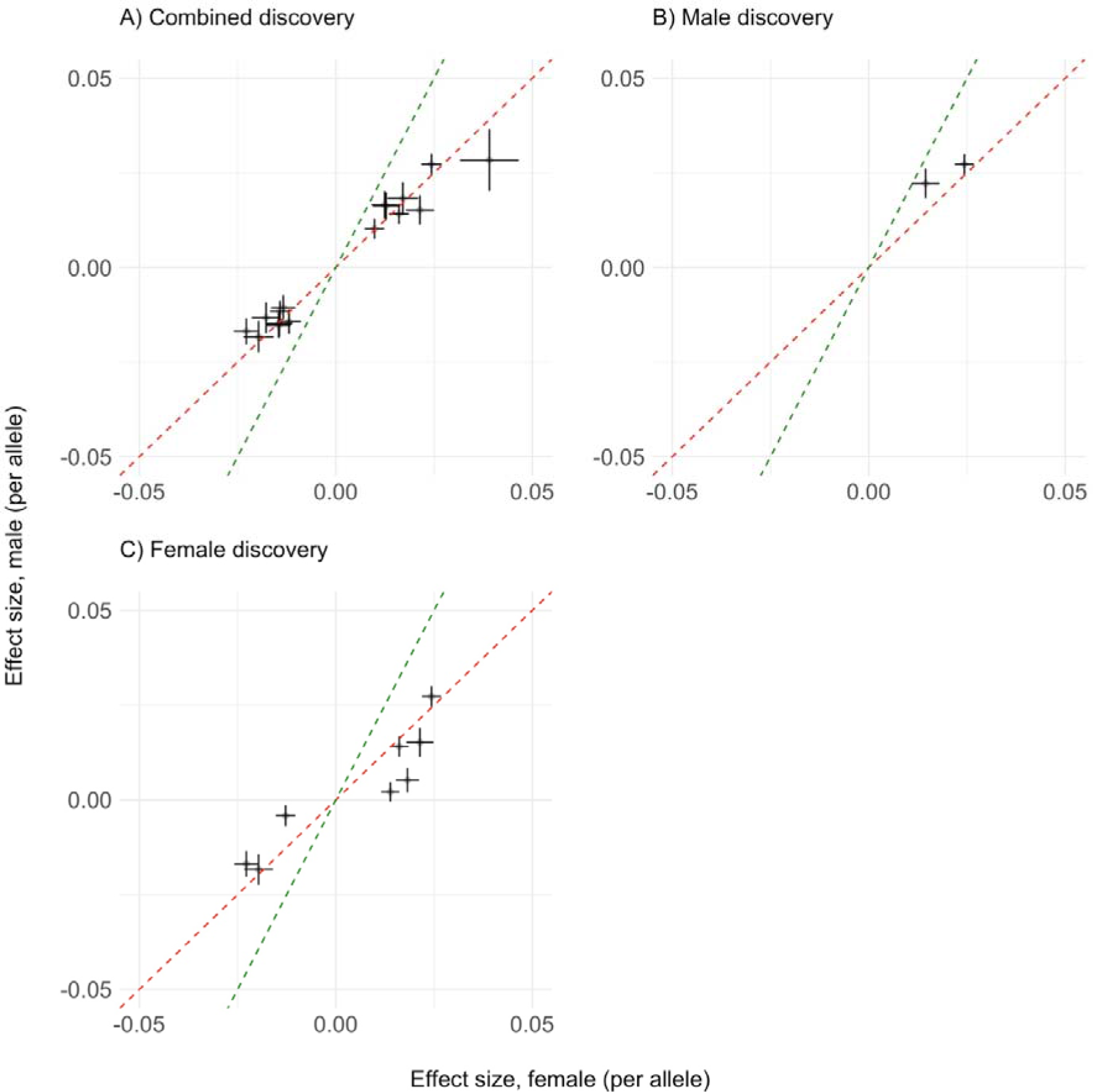
Comparison of per-allele effects from sex-specific analyses (+/− SE) of lead SNPs in PAR as identified in a **A)**combined discovery set (N=16), **B)**male discovery set (N=2) or **C)**female discovery set (N=8). The green and red dashed lines indicate the expectations under full DC and escape from X-inactivation, respectively. DCC was not estimated due to low number of lead SNPs in PAR.

**Supplementary Figure 5.**
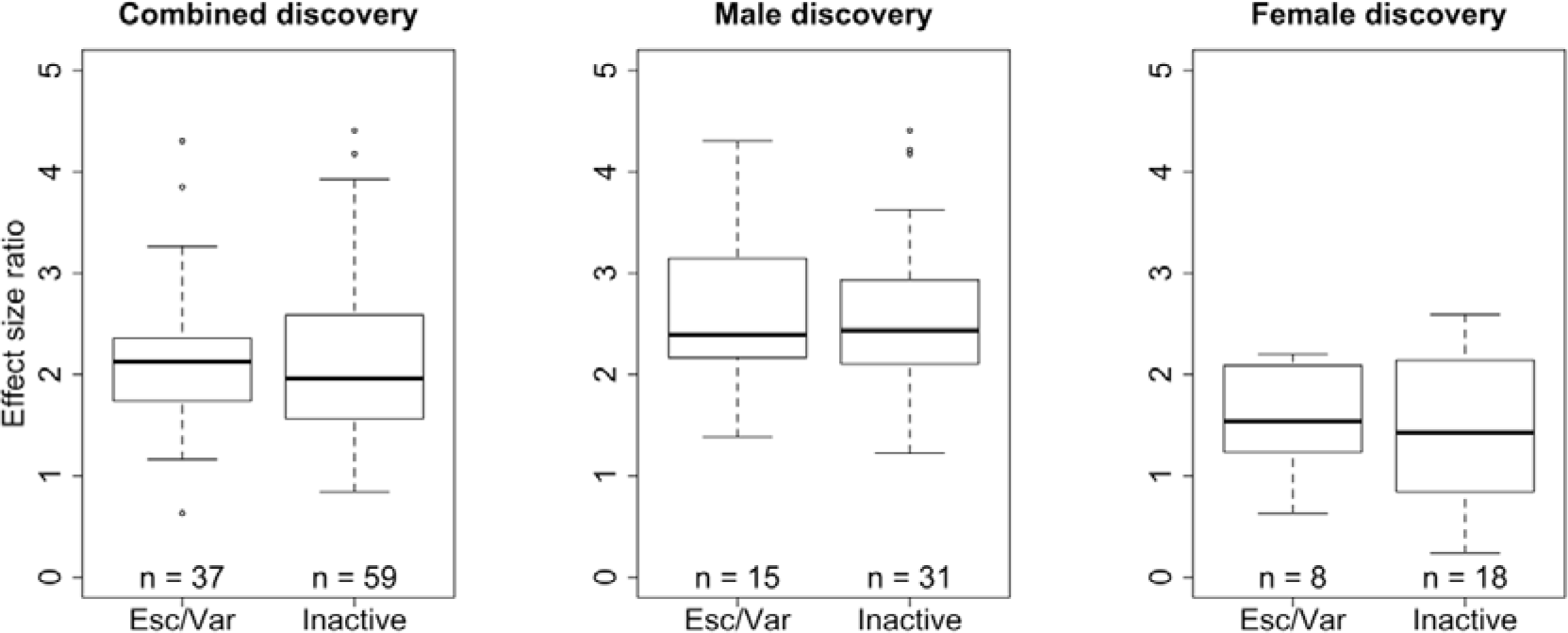
Effects size ratios for the lead SNPs across analysed complex traits are compared between “Escape/Variable” and “Inactive” groups, which include SNPs physically located within a gene region with previously reported XCI status (Tukiainen, A.-C. Villani, *et al.*, 2017). We exclude variants in the regions of heterogeneity as well as 2 variants with the absolute ratio values > 10 (male discovery sample).

**Supplementary Figure 6.**
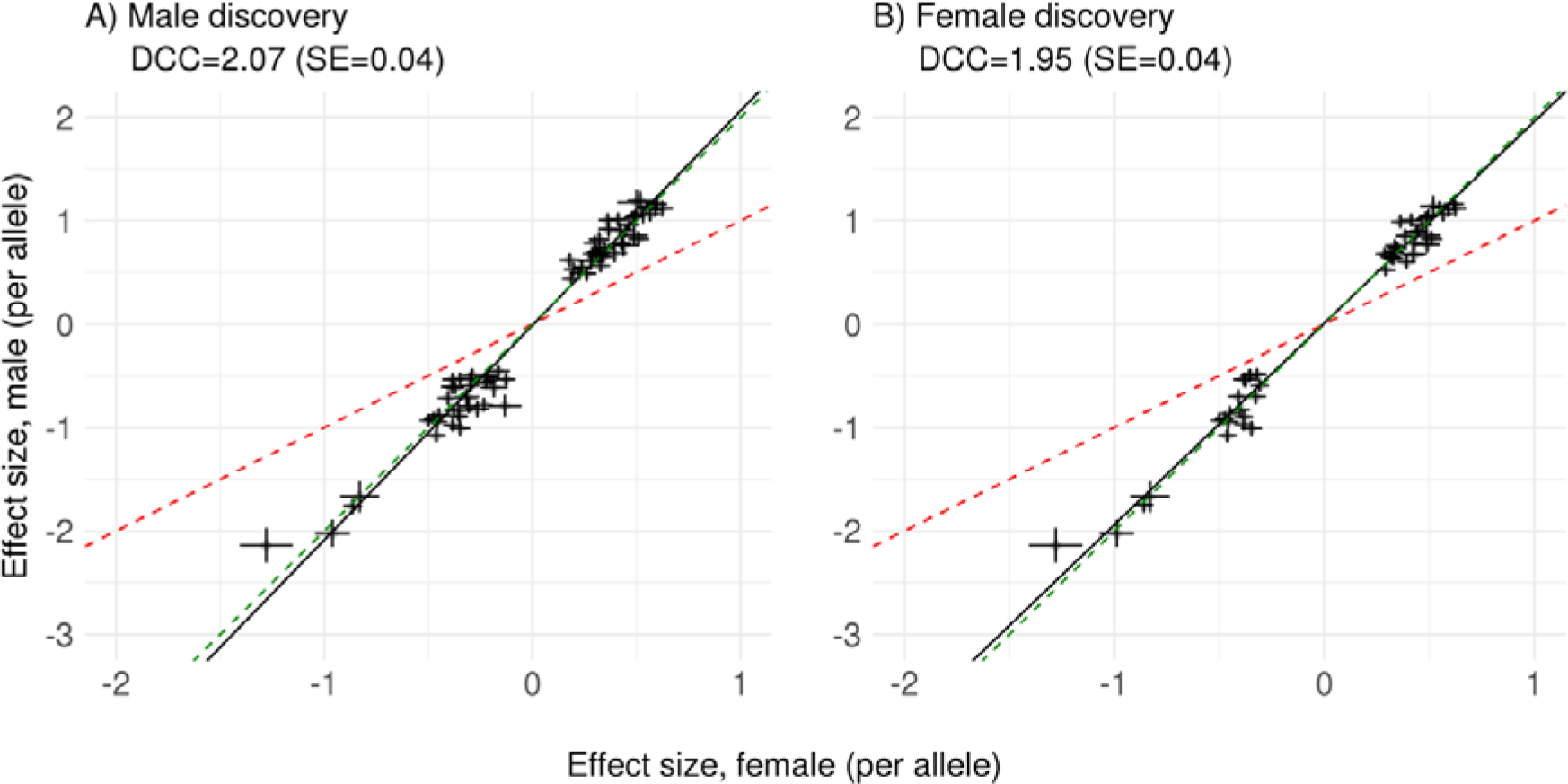
Comparison of per-allele effects from sex-specific analyses (+/− SE) for X-chromosome *cis*-eQTLs in CAGE whole blood. DCC of 1.95 (SE=0.04) is observed for 51 eQTLs (P<1.6×10^−10^) in the female discovery analysis, and DCC of 2.07 (SE=0.04) for 74 eQTLs (P<1.6×10^−10^) in the male discovery analysis. The green and red dashed lines indicate the expectations under full DC and escape from X-inactivation, respectively. The black line represents DCC.

**Supplementary Figure 7.**
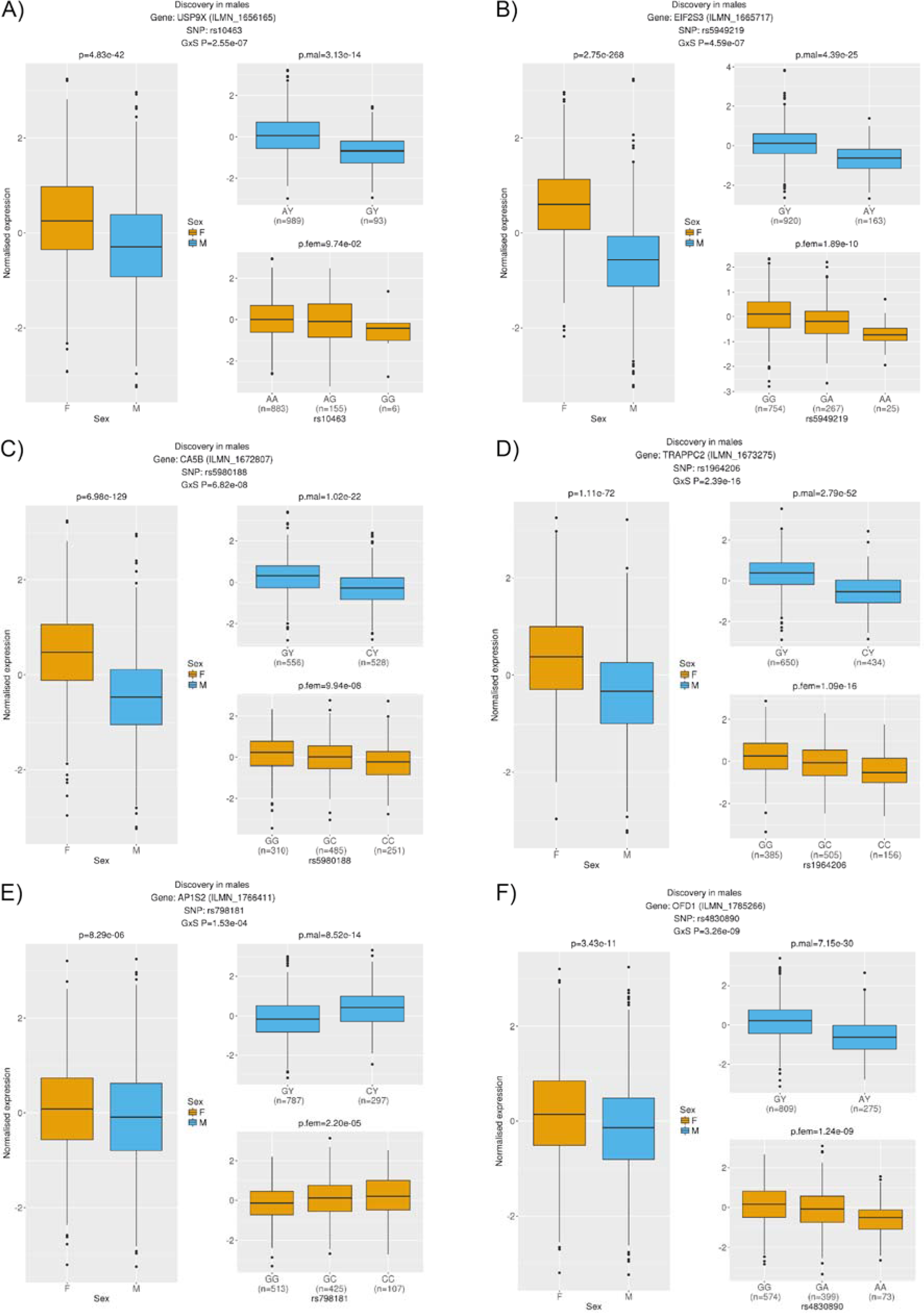
A total of 6 eQTLs identified in the male discovery *cis*-eQTL analysis in CAGE whole blood are annotated to escape XCI. These genes show higher expression in females compared to males (P<3.1×10^−3^, i.e. 0.05/16), as expected for genes that escape from XCI, but also significant differences between the effect estimate of the top associated SNP on gene expression after correction for mean differences in expression between the sexes (genotype-by-sex interaction P<3.1×10^−3^), which is consistent with FDC. This suggests that sexual dimorphism in these genes may not be due to escape from XCI. Orange corresponds to females. Blue corresponds to males.

**Supplementary Figure 8.**
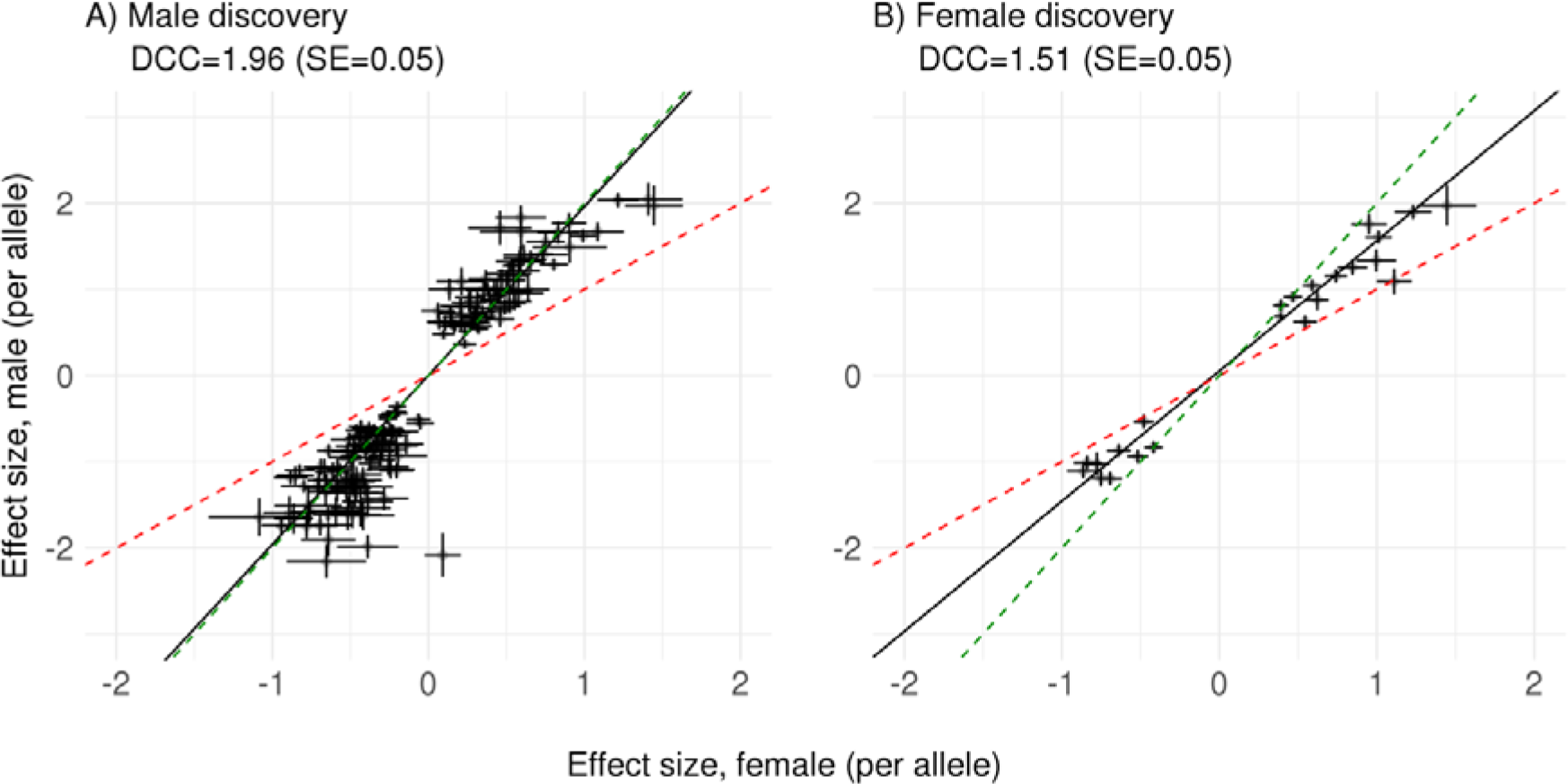
The per-allele effect estimates of top eQTLs across all 22 tissues in GTEx in the discovery sex is compared to the corresponding eQTL in the other sex from the matching tissue. DCC of 1.96 (SE=0.05) is observed for 175 eQTLs in the male discovery analysis, and 1.51 (SE=0.05) for 23 eQTLs in the female discovery analysis. The green and red dashed lines indicate the expectations under full DC and escape from X-inactivation, respectively. The black line represents DCC.

**Supplementary Figure 9.**
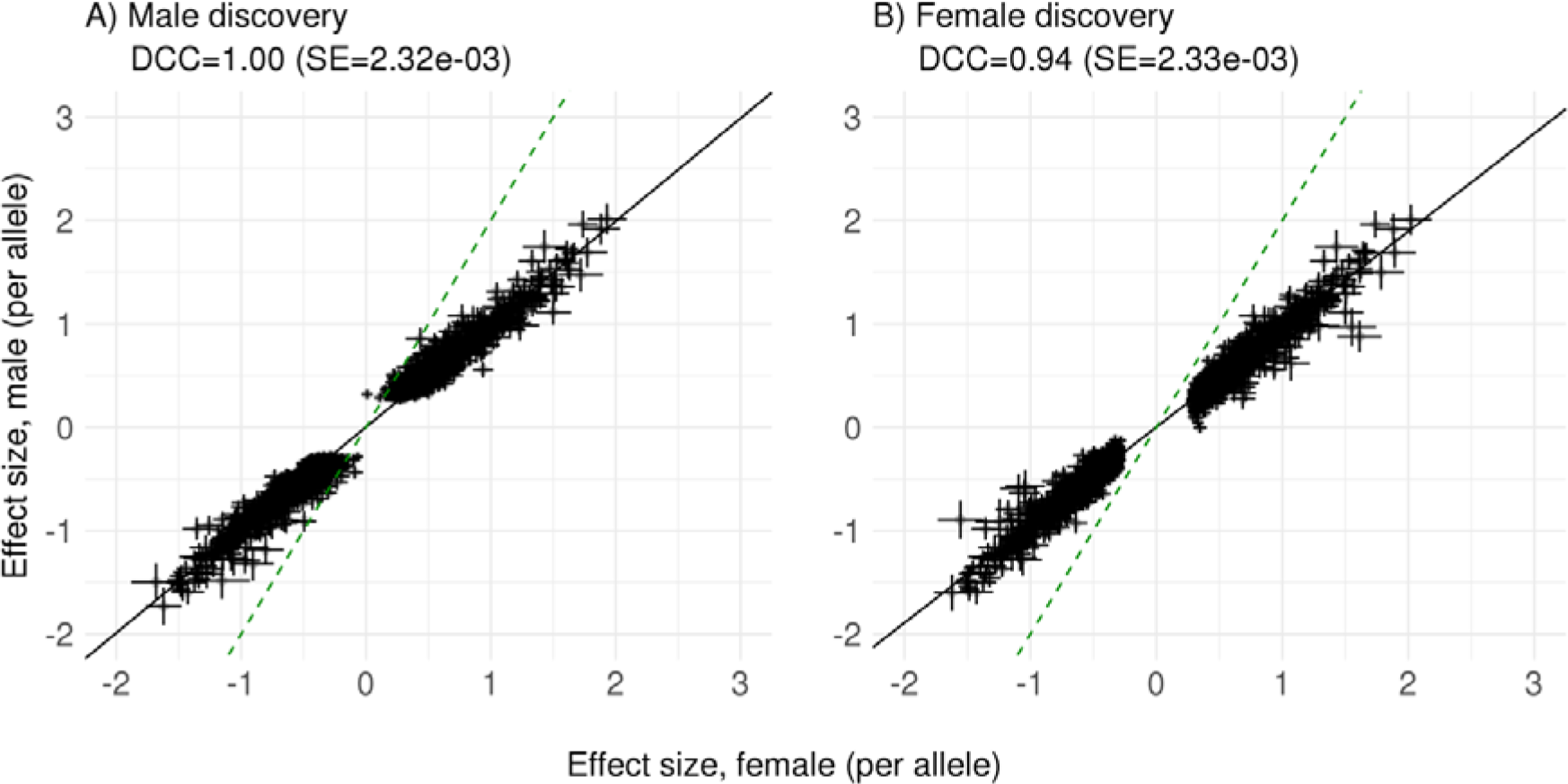
Comparison of per-allele effects from sex-specific analyses (+/− SE) for autosomal *cis*-eQTLs identified in CAGE whole blood. DCC is expected to be equal in males and females. DCC of 1.00 (SE=2.3×10^−3^) is observed for 3,116 eQTLs with P<10^−10^ in the male discovery analysis, and 0.94 (SE=2.3×10^−3^) for 3,165 eQTLs with P<10^−10^ in the female discovery analysis. The green dashed line represents the *y*=2*x* line. The black line represents DCC.

**Supplementary Figure 10.**
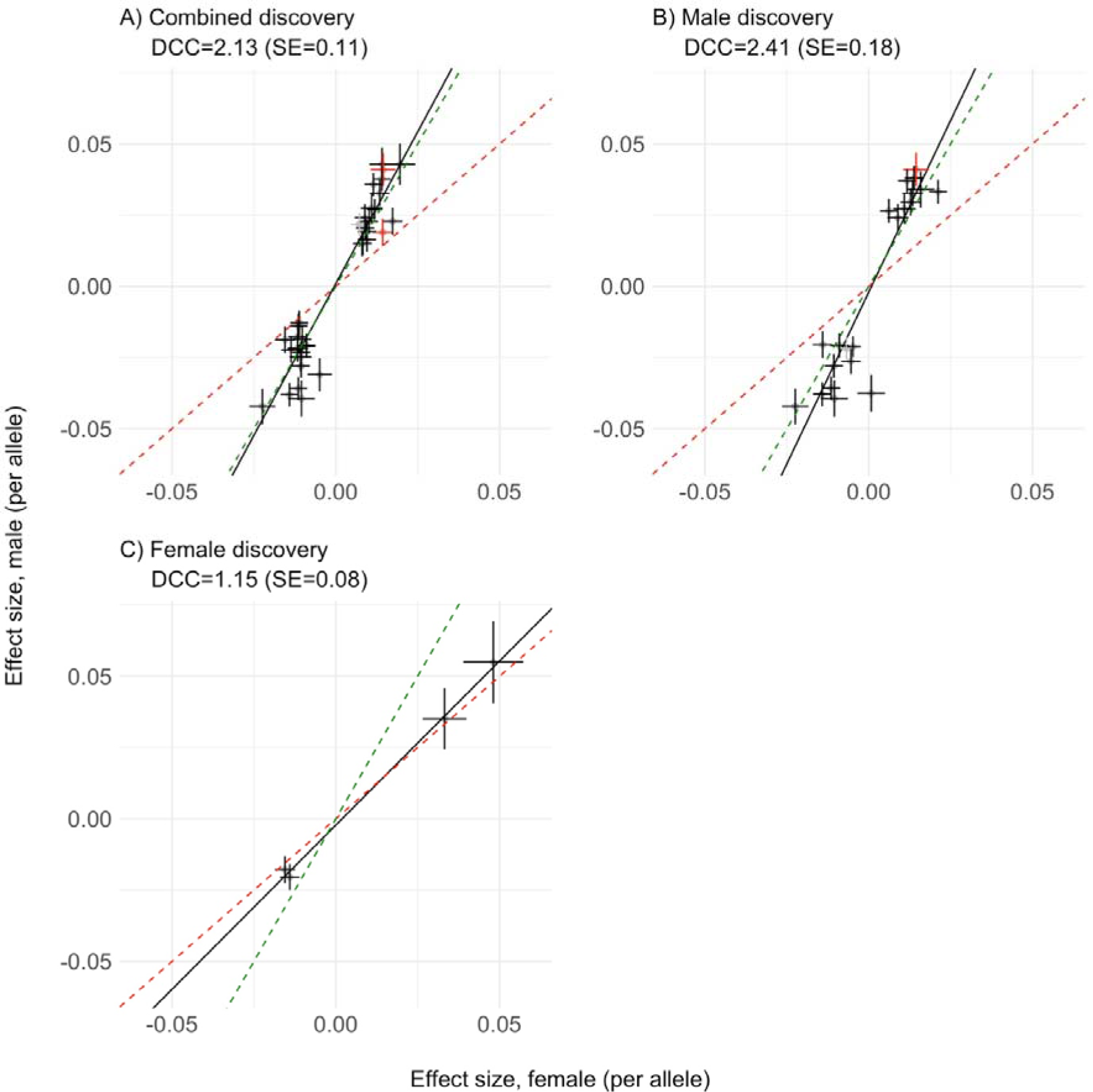
Comparison of per-allele effects from sex-specific analyses (+/− SE) of the SNPs associated with complex traits through gene expression, as identified in a A) combined male-female SMR analysis (N=37), and sex-stratified SMR analyses (B, N=23; C, N=4). The SNPs are coloured according to the reported inactivation status of the genes that showed evidence of pleiotropic association with phenotypic traits (SMR genes, red “Escape/Variable”, black = “Inactive”, grey = “Unknown”). The results are presented in the **Supplementary Tables 12-14**. The green and red dashed lines indicate the expectations under full DC and escape from X-inactivation, respectively. The black line represents DCC.

## Supplementary Tables

Supplementary Tables 7-9, 11-18 are provided as Excel spreadsheets.

**Supplementary Table 1a.**
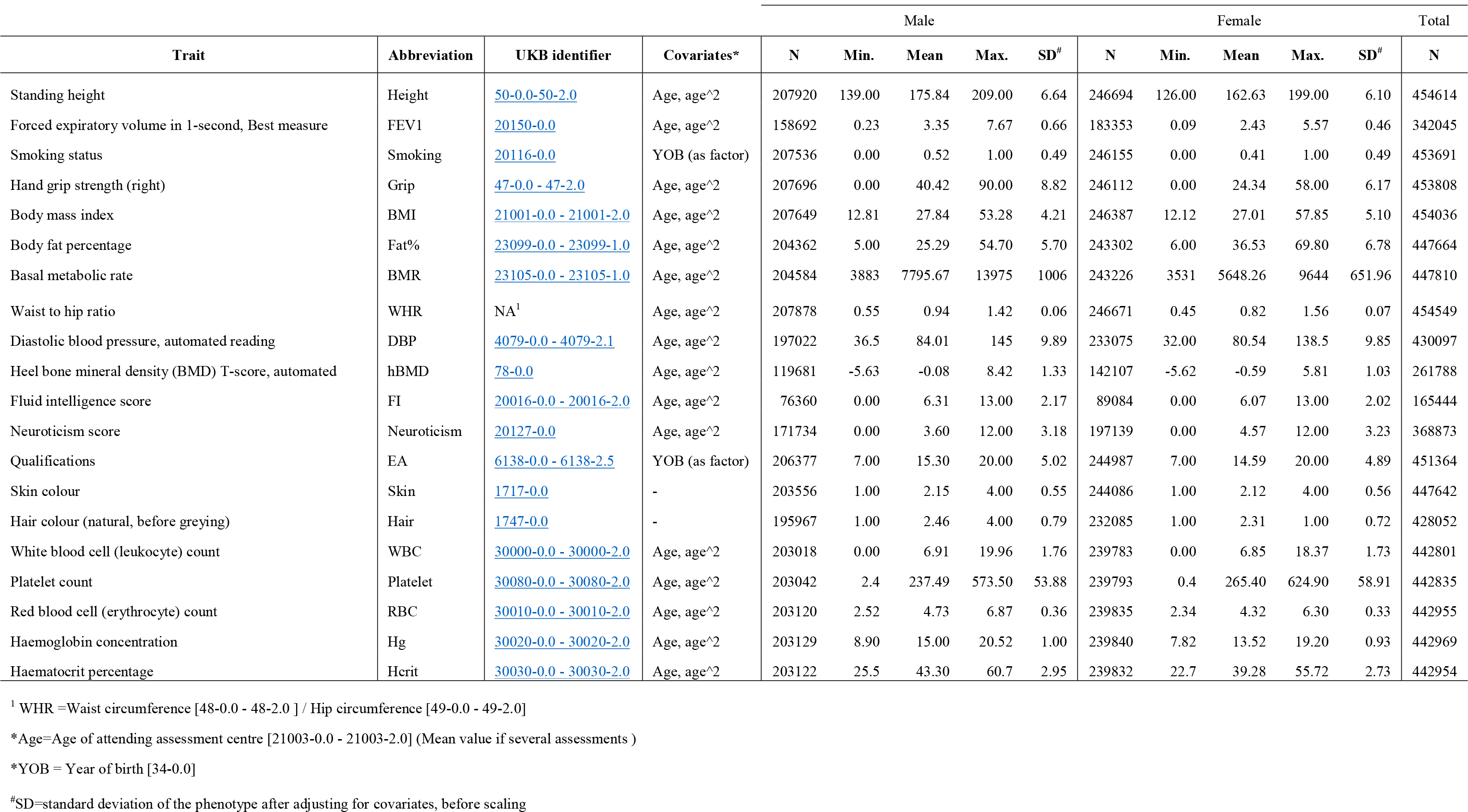
A) The UK Biobank trait information

**Supplementary Table 1b.**
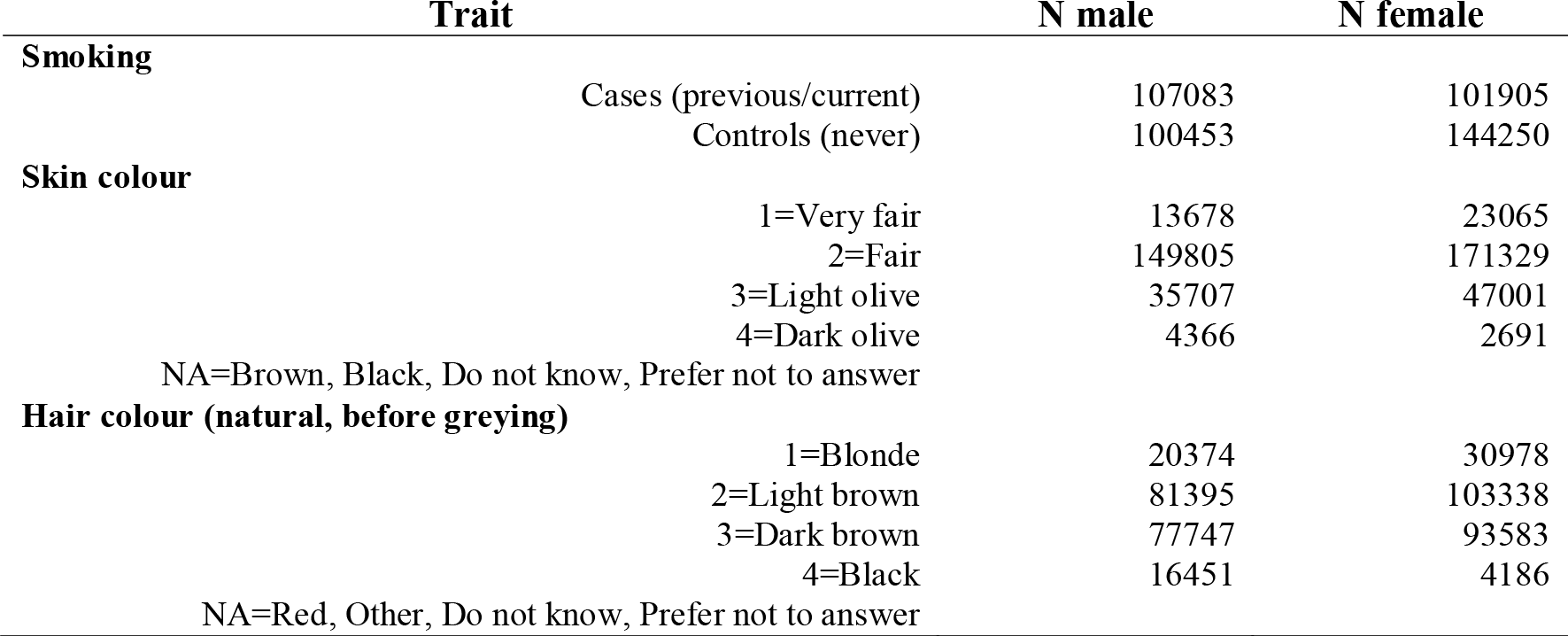
B) The UK Biobank trait information

**Supplementary Table 2.** The X-chromosome-wide (non-PAR) heritability estimates and DC ratio estimates obtained with REML analysis and estimated from GWAS summary statistics.

| REML summary statistics |  |  |  |  |  |  |
| --- | --- | --- | --- | --- | --- | --- |
| Trait | N<sub>m</sub> | N<sub>f</sub> | $h^2_{\text{SNP, M}}$<br>(SE, %) | $h^2_{\text{SNP, F}}$<br>(SE, %) | $h^2_{\text{SNP, M}}/h^2_{\text{SNP, F}}$<br>(SE) | DC ratio<br>(SE) |
| **Height** | 99762 | 99796 | 1.55 (0.12) | 0.88 (0.09) | 1.76 (0.23) | 1.59 (0.07) |
| **FEV1** | 91543 | 89326 | 0.57 (0.1) | 0.36 (0.09) | 1.60 (0.47) | 1.35 (0.16) |
| **Smoking** | 99566 | 99584 | 0.48 (0.09) | 0.18 (0.07) | 2.67 (1.19) | 1.98 (0.42) |
| **Grip** | 99651 | 99551 | 0.42 (0.08) | 0.28 (0.07) | 1.52 (0.51) | 1.78 (0.21) |
| **BMI** | 99634 | 99663 | 0.97 (0.11) | 0.42 (0.08) | 2.28 (0.52) | 2.13 (0.21) |
| **Fat%** | 98070 | 98362 | 0.97 (0.11) | 0.38 (0.08) | 2.53 (0.62) | 2.85 (0.32) |
| **BMR** | 98162 | 98321 | 1.22 (0.12) | 0.58 (0.09) | 2.12 (0.40) | 2.54 (0.19) |
| **WHR** | 99727 | 99798 | 0.53 (0.09) | 0.16 (0.07) | 3.30 (1.48) | 1.84 (0.25) |
| **DBP** | 94565 | 94166 | 0.27 (0.08) | 0.31 (0.08) | 0.85 (0.33) | 0.94 (0.19) |
| **hBMD** | 90779 | 90800 | 0.52 (0.09) | 0.27 (0.07) | 1.95 (0.64) | 2.13 (0.33) |
| **FI** | 59641 | 59650 | 0.57 (0.13) | 0.45 (0.12) | 1.25 (0.45) | 1.13 (0.27) |
| **Neuroticism** | 98925 | 95683 | 0.38 (0.08) | 0.17 (0.07) | 2.19 (1.00) | 1.51 (0.34) |
| **EA** | 99023 | 99147 | 0.33 (0.08) | 0.45 (0.09) | 0.72 (0.22) | 0.82 (0.13) |
| **Skin** | 97746 | 98743 | 0.2 (0.07) | 0.03 (0.06) | 5.82 (9.46) | 1.95 (0.74) |
| **Hair** | 94080 | 93995 | 0.23 (0.07) | 0.02 (0.05) | 10.02 (23.45) | 2.05 (0.76) |
| **WBC** | 97422 | 96987 | 0.52 (0.09) | 0.17 (0.07) | 3.07 (1.36) | 2.53 (0.46) |
| **Platelet** | 97426 | 96991 | 0.55 (0.09) | 0.19 (0.07) | 2.90 (1.10) | 2.55 (0.38) |
| **RBC** | 97460 | 97009 | 0.61 (0.1) | 0.26 (0.07) | 2.36 (0.77) | 2.46 (0.34) |
| **Hgb** | 97474 | 97006 | 0.77 (0.11) | 0.26 (0.08) | 2.91 (0.92) | 5.22 (0.77) |
| **Hcrit** | 97466 | 97007 | 0.74 (0.11) | 0.16 (0.07) | 4.51 (2.00) | 5.07 (0.83) |
| Mean (SD) | 95406 | 95079 | 0.62 (0.34) | 0.30 (0.20) | 2.82 (2.07) | 2.22 (1.14) |

**Supplementary Table 3.** Estimated DC ratios and genetic correlations (r_g_) on the X chromosome and autosomes.

|  | **X chromosome** |  | **Autosomes** |  |
| --- | --- | --- | --- | --- |
|  | **DC** | **$r_g$** | **DC** | **$r_g$** |
| Height | 1.59 (0.07) | 0.96 (0.009) | 0.98 (0.01) | 0.96 (0.001) |
| FEV1 | 1.35 (0.16) | 0.93 (0.03) | 1.01 (0.02) | 0.96 (0.003) |
| Smoking | 1.98 (0.42) | 0.95 (0.059) | 1.04 (0.03) | 0.85 (0.005) |
| Grip | 1.78 (0.21) | 0.82 (0.031) | 1.17 (0.03) | 0.86 (0.004) |
| BMI | 2.13 (0.21) | 0.80 (0.03) | 1.02 (0.02) | 0.94 (0.002) |
| Fat% | 2.85 (0.32) | 0.57 (0.053) | 1.00 (0.02) | 0.89 (0.002) |
| BMR | 2.54 (0.19) | 0.92 (0.018) | 1.13 (0.01) | 0.94 (0.002) |
| WHR | 1.84 (0.25) | 0.75 (0.039) | 0.83 (0.02) | 0.72 (0.004) |
| DBP | 0.94 (0.19) | 0.74 (0.057) | 0.67 (0.02) | 0.92 (0.004) |
| hBMD | 2.13 (0.33) | 0.97 (0.043) | 0.66 (0.01) | 0.91 (0.004) |
| FI | 1.13 (0.27) | 0.81 (0.069) | 0.96 (0.03) | 1.00 (0.006) |
| Neuroticism | 1.51 (0.34) | 0.94 (0.067) | 0.94 (0.03) | 0.90 (0.006) |
| EA | 0.82 (0.13) | 0.94 (0.04) | 0.94 (0.02) | 0.93 (0.004) |
| Skin | 1.95 (0.74) | 0.81 (0.12) | 0.84 (0.02) | 0.98 (0.003) |
| Hair | 2.05 (0.76) | 0.72 (0.129) | 0.92 (0.01) | 0.99 (0.002) |
| WBC | 2.53 (0.46) | 0.76 (0.053) | 0.9 (0.01) | 0.96 (0.003) |
| Platelet | 2.55 (0.38) | 0.78 (0.04) | 0.91 (0.01) | 0.96 (0.002) |
| RBC | 2.46 (0.34) | 0.64 (0.068) | 0.94 (0.01) | 0.93 (0.003) |
| Hgb | 5.22 (0.77) | 0.65 (0.044) | 1.03 (0.02) | 0.91 (0.004) |
| Hcrit | 5.07 (0.83) | 0.51 (0.05) | 1.03 (0.02) | 0.91 (0.004) |

**Supplementary Table 4.** Regions of heterogeneity.

| Region | Top SNP | Top-SNP bp | Heterogeneity P-value | Left Bound (bp) | Right Bound( bp) | Span (kb) | Traits |
| --- | --- | --- | --- | --- | --- | --- | --- |
| **Region 1** | rs17307280 | 8916646 | 4.34E-12 | 8635709 | 8929104 | 293 | hBMD |
| **Region 1** | rs112265145 | 8906893 | 2.10E-44 | 8635709 | 8929104 | 293 | Hgb |
| **Region 1** | rs56066690 | 8912070 | 8.45E-46 | 8635709 | 8929104 | 293 | Hcrit |
| **Region 1** | rs56066690 | 8912070 | 1.83E-27 | 8635709 | 8929104 | 293 | RBC |
| **Region 1** | rs745535498 | 8912871 | 6.62E-09 | 8635709 | 8929104 | 293 | Fat% |
| **Region 2** | rs12556728 | 30402866 | 2.61E-08 | 30320507 | 30572217 | 251 | Height |
| **Region 3** | rs56908677 | 56958534 | 4.97E-09 | 55058361 | 65331684 | 10273 | Hgb |
| **Region 3** | rs56908677 | 56958534 | 6.93E-09 | 55058361 | 65331684 | 10273 | Hcrit |
| **Region 4** | rs113121621 | 66389189 | 4.00E-08 | 56197395 | 67837267 | 11639 | Fat% |

**Supplementary Table 5.** Estimates of dosage compensation (DC) and genetic correlation (r_g_) after excluding regions of heterogeneity. The DC and r_g_ are marked as follows: 0 - including all SNPs, 1- excluding the SNPs in the region 1; 2- excluding the SNPs in the region 2; 3- excluding the SNPs in the region 3; 4- excluding the SNPs in the region 4; 13- excluding the SNPs in the region 1 and region 3; 14- excluding the SNPs in the region 1 and region 4

|  | DC0 | Rg0 | DC1 | Rg1 | DC2 | Rg2 | DC3 | Rg3 | DC4 | Rg4 | DC13 | Rg13 | DC14 | Rg14 |
| --- | --- | --- | --- | --- | --- | --- | --- | --- | --- | --- | --- | --- | --- | --- |
| **hBMD** | 2.13<br>(0.33) | 0.97<br>(0.043) | 2.08<br>(0.33) | 0.98<br>(0.02) | - | - | - | - | - | - | - | - | - | - |
| **Fat%** | 2.85<br>(0.32) | 0.57<br>(0.053) | 2.81<br>(0.32) | 0.57<br>(0.03) | - | - | - | - | 2.21<br>(0.26) | 0.74<br>(0.02) | - | - | 2.16<br>(0.26) | 0.74<br>(0.02) |
| **Hgb** | 5.22<br>(0.77) | 0.65<br>(0.044) | 4.87<br>(0.73) | 0.68<br>(0.02) | - | - | 2.54<br>(0.42) | 0.68<br>(0.02) | - | - | 2.15<br>(0.37) | 0.74<br>(0.02) | - | - |
| **Hcrit** | 5.07<br>(0.83) | 0.51<br>(0.05) | 4.66<br>(0.76) | 0.53<br>(0.02) | - | - | 2.53<br>(0.43) | 0.62<br>(0.03) | - | - | 2.12<br>(0.38) | 0.68<br>(0.03) | - | - |
| **Height** | 1.59<br>(0.07) | 0.96<br>(0.009) | - | - | 1.58<br>(0.07) | 0.97<br>(0.003) | - | - | - | - | - | - | - | - |
| **RBC** | 2.46<br>(0.34) | 0.64<br>(0.068) | 2.27<br>(0.32) | 0.67<br>(0.04) | - | - | - | - | - | - | - | - | - | - |

**Supplementary Table 6.** Number of lead SNPs identified in sex-stratified and combined analyses (GCTA-COJO). The number of SNPs retained after exclusion of markers located in the regions of male-female heterogeneity for six traits is indicated parentheses.

|  | Male discovery |  | Female discovery |  | Combined discovery |  |
| --- | --- | --- | --- | --- | --- | --- |
|  | Non-PAR | PAR | Non-PAR | PAR | Non-PAR | PAR |
| BMI | 10 | -- | 2 | -- | 19 | 1 |
| BMR | 23 | -- | 4 | -- | 37 | 1 |
| DBP | 0 | -- | 1 | -- | 1 | -- |
| EA | 1 | -- | 0 | -- | 5 | -- |
| Fat% | 10 (7) | -- | 2 | -- | 16 (13) | 1 |
| FEV1 | 5 | -- | 3 | -- | 14 | -- |
| Grip | 6 | -- | 1 | -- | 10 | -- |
| Hair | 1 | -- | 1 | -- | 5 | -- |
| Hcrit | 8 (6) | -- | 4 | -- | 15 (13) | -- |
| hBMD | 4 (3) | -- | 2 | -- | 5 (4) | 1 |
| Height | 46 (45) | 2 | 24 | 7 | 64 (63) | 11 |
| Hgb | 6 (4) | -- | 3 | -- | 13 (11) | -- |
| Neuroticism | 0 | -- | 0 | -- | 3 | -- |
| Platelet | 13 | -- | 8 | -- | 15 | -- |
| RBC | 9 (8) | -- | 3 | 1 | 16 (15) | 1 |
| Skin | 1 | -- | 0 | -- | 1 | -- |
| Smoking | 3 | -- | 1 | -- | 3 | -- |
| WBC | 5 | -- | 1 | -- | 9 | -- |
| WHR | 2 | -- | 2 | -- | 10 | -- |
| Total | 153 (143) | 2 | 62 | 8 | 261 (251) | 16 |

**Supplementary Table 10.** Samples size and number of X-linked transcripts expressed per tissue-type in GTEx. X-chromosome *cis*-eQTL analysis is performed in 22 tissue samples for which within tissue sample size was greater than N=50 in both males and females.

| **Tissue** | **Total** | **No. Females** | **No. Males** | **No. transcripts** |
| --- | --- | --- | --- | --- |
| Colon Sigmoid | 124 | 50 | 74 | 792 |
| Adrenal Gland | 126 | 56 | 70 | 773 |
| Pancreas | 149 | 62 | 87 | 726 |
| Heart Atrial Appendage | 159 | 54 | 105 | 766 |
| Colon Transverse | 169 | 72 | 97 | 808 |
| Stomach | 170 | 72 | 98 | 802 |
| Breast Mammary Tissue | 183 | 80 | 103 | 863 |
| Adipose Visceral Omentum | 185 | 67 | 118 | 802 |
| Heart Left Ventricle | 190 | 67 | 123 | 730 |
| Skin Not Sun Exposed Suprapubic | 196 | 67 | 129 | 818 |
| Artery Aorta | 197 | 71 | 126 | 832 |
| Esophagus Muscularis | 218 | 81 | 137 | 806 |
| Esophagus Mucosa | 241 | 90 | 151 | 793 |
| Nerve Tibial | 256 | 93 | 163 | 896 |
| Cells Transformed fibroblasts | 272 | 102 | 170 | 733 |
| Lung | 278 | 96 | 182 | 907 |
| Thyroid | 278 | 99 | 179 | 916 |
| Artery Tibial | 285 | 101 | 184 | 833 |
| Adipose Subcutaneous | 298 | 111 | 187 | 855 |
| Skin Sun Exposed Lower leg | 302 | 112 | 190 | 854 |
| Whole Blood | 338 | 125 | 213 | 729 |
| Muscle Skeletal | 361 | 133 | 228 | 745 |

## Methods and Materials

### Genotype coding

The summary statistics reported in this study were generated with a combination of BOLT-LMM v2.3 (Loh *et al.*, 2018), GCTA 1.94 (Yang, Lee, *et al.*, 2011), and PLINK 1.90 (Purcell *et al.*, 2007), all of which have default settings for the treatment of X-chromosome SNPs. For analyses performed using PLINK, we used the default parameters which codes males as {0,1}, and thus gives the appropriate per-allele effect estimates. For BOLT-LMM and GCTA, the male genotypes were analysed as diploid using a {0,2} coding. This distinction makes no impact on the strength of association (i.e. P-values), however, we multiply the effect estimates and the corresponding standard errors from the diploid male-specific analysis by 2, allowing us to report our results as per-allele effect estimates. In all cases, females were coded as {0,1,2}.

### Data

#### UK Biobank data

Sex-stratified association analyses of 20 complex was performed using the phenotype data on N_m_=208,419 males and N_f_=247,186 females of European-ancestry and UKB Version 3 release of imputed genotype data (6,871 SNPs in pseudoautosomal region (PAR) and 253,842 SNPs in non-pseudoautosomal region (non-PAR) that satisfied our quality control criteria and had minor allele frequency, MAF>0.01). The phenotypes were adjusted for appropriate covariates and converted to sex-specific Z-scores prior to analysis (See **Supplementary Table 1**and Supplementary Methods and Material for full details).

#### CAGE gene expression data

Gene expression and X-chromosome genotype data were available in a subset of N=2,130 individuals of verified European ancestry (N_m_=1,084 males, N_f_=1,046 females) from the Consortium for the Architecture of Gene Expression (CAGE) (Lloyd-jones *et al.*, 2017). A total of 36,267 autosomal and 1,639 X-chromosome gene expression probes (28 in the PAR) in whole-blood were available for analysis following quality control. Gene expression levels were adjusted for PEER factors (Stegle *et al.*, 2010, 2012) that were not associated with sex (P_sex_>0.05) in order to preserve the effect of sex on expression and where available, measured covariates such as age, cell counts, and batch effects. A total of 1,066,905 HapMap3 SNPs imputed to 1000 Genomes Phase 1 Version 3 reference panel (Altshuler *et al.*, 2012) and 190,245 non-PAR X-chromosome SNPs (minor allele frequency, MAF>0.01) imputed to the Haplotype Reference Consortium (HRC, release 1.1) (McCarthy *et al.*, 2016) were available for analysis.

#### GTEx gene expression data

We used the fully-processed, normalised and filtered RNA-seq data from the Genotype Tissue Expression project (GTEx v6p release). X-chromosome imputed SNP data was obtained from dbGap (Accession phs000424.v6.p1). We restricted our analyses to 22 tissue samples for which within tissue sample size was greater than N=50 in both males and females (**Supplementary Table 10**). A total of 1,121 transcripts (31 in the PAR) were expressed in at least one tissue, with a mean of 808 transcripts expressed across all 22 tissues (**Supplementary Table 10**) and a total of 127,808 imputed SNPs in the non-PAR of the X chromosome (MAF>0.05).

### Statistical analysis

#### Sex-stratified XWAS

Summary statistics were generated for 20 complex traits in the UK Biobank using BOLT-LMM v2.3 (Loh *et al.*, 2018) for the X-chromosome SNPs with MAF>0.01 in both sexes and using 561,572 HapMap3 SNPs (autosomal and X-chromosomal, pairwise R^2^ <0.9) as “model SNPs” to estimate genetic relationship matrix (GRM) and correct for confounding.

#### Combined analyses

For complex traits, the results from the sex-stratified association testing were meta-analysed using the inverse-variance weighted method to obtain combined results (performed in R). For combined analysis of gene expression traits, individual data from males and female were pooled together. We assumed full DC for all loci for these analyses.

#### Significant SNP-trait associations

GCTA-COJO (Yang *et al.*, 2012) was used to identify sets of jointly significant SNPs associated with a trait at genome-wide significance (GWS) threshold P<5.0×10^−8^. We use genotypes of a random sample of 100,000 unrelated UKB females of European ancestry as a linkage disequilibrium (LD) reference and increase a distance of assumed complete linkage equilibrium between markers (window size) to 50Mb due to higher levels of LD on the X chromosome.

#### Estimation of dosage compensation ratio and genetic correlation from summary statistics

Following (Lee *et al.*, 2018), we calculated the DC ratio for 20 complex traits from the summary statistics of the sex-stratified X-chromosome analysis using the following equation:

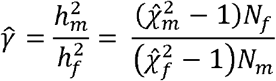

Where 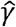,is the estimate of the DCR; 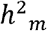 and 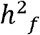 are the M/F SNP-heritabilities; 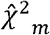 and 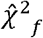 are the mean chi-square estimates from association analysis; and *N*_*m*_ and, *N*_*f*_ are the corresponding sample sizes used in the analysis, respectively.

The corresponding standard error is estimated as:

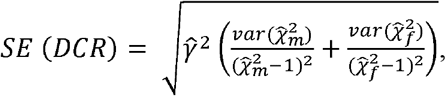

where the 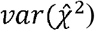 is the variance of the mean test statistic across the X chromosome, which is approximately equal to 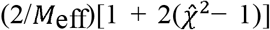 *M*_*eff*_ is the effective number of SNPs, which for the X chromosome is approximately equal to 1,300 (Lee *et al.*, 2018). The DC ratio of 2 indicates the evidence for full DC, while the value of 0.5 implies complete escape from inactivation (no DC).

We also we obtained an estimator for the male-female genetic correlation on the X chromosome (non-PAR region) or autosomes using the following equation,

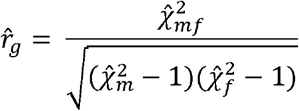

where, as before, 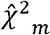 and 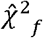 are the mean chi-square estimates from association analysis and 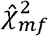 is the cross-product of the Z-statistics from the male and female analyses.

We calculate standard errors using a block jackknife method. We assign SNPs across the X chromosome to blocks (B=1000) and for each block *k* we calculate an estimate of the genetic correlation 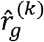 as above excluding the SNPs in this block. The standard error is then calculated as follows:

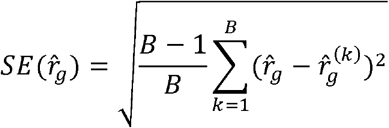

#### Heterogeneity in SNP effects on complex traits

To test the difference in the SNP effects estimated in male or female datasets we apply a heterogeneity test. If 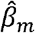 and 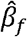 are the male and female per-allele effect estimates, and 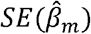 and 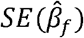 are their corresponding standard errors, then we used the test statistic

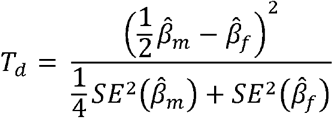

which follows a χ^2^-distribution with one degree of freedom under the null hypothesis of no difference in estimates under full DC assumption. We set a P-value threshold of P<5.0×10^−8^ to identify the markers with significant difference in estimated effects and further apply LD-clumping (R^2^ threshold 0.05) to identify regions of heterogeneity. The coordinates of protein coding genes in these regions were extracted with BioMart tool (See URLs), using the genome assembly GRCh37.p13 from Genome Reference Consortium.

### Estimation of the SNP-heritability

We estimated the proportion of variance explained by X-chromosome SNPs in males and females separately using GREML and a genome partitioning approach as in (Yang, Manolio, *et al.*, 2011), which is implemented in the GCTA software package (Yang, Lee, *et al.*, 2011). Here, we model the trait as,

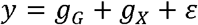

where, *y* is a N x 1 vector of phenotype for each trait, with sample size N; gG is an N × 1 vector of the total genetic effects from the autosome with 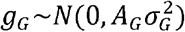 where *A*_*G*_ is the GRM between individuals estimated from 548,860 autosomal HapMap3 SNPs; *g*_*X*_ is an N × 1 vector of X-linked genetic effects with 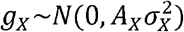, where *A*_*X*_ is a GRM calculated from 253,842 X-chromosome SNPs; and 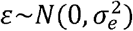 is the residual. Partitioning in this way will allow for an estimation of the parameter 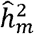 conditional on the autosomal GRM. Thus, we can estimate the proportion of phenotypic variance that is due to the X chromosome while controlling for sample structure captured by genetic variants on the autosome (Yang, Manolio, *et al.*, 2011). We applied this model to the 20 complex traits, limiting our analysis to a maximum of 100,000 unrelated males or females due to computational restrictions.

The standard errors of the M/F ratio of the estimated SNP-heritabilities on the X chromosome was estimated as,

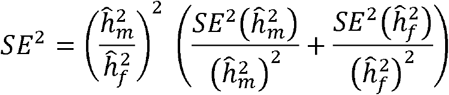

where 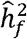 and 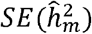 are the GREML-estimates of SNP-heritability in males and females, respectively, and 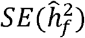 and 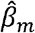 are the corresponding standard errors.

#### Sex-stratified X-chromosome and autosomal *cis*-eQTL analysis

Gene expression levels were modelled as a linear function of the number of reference alleles for SNPs on the same chromosome in males and females, separately. We used GCTA and PLINK to analyse the CAGE and GTEx datasets, respectively. Sample structure was accounted for by adjusting for genotyping principal components and PEER factor in the GTEx analysis, and a random polygenic effect captured by an autosomal genetic relationship matrix in the CAGE analysis. For each gene expression probe/transcript, we identified the top associated SNP that satisfied a Bonferroni corrected significance threshold in the discovery sex (i.e. eQTL), and extracted the same eQTL in the other sex to compare the per-allele eQTL effect estimates between the sexes (see **Estimating effect size ratio and dosage compensation coefficient, below**).

#### Summary data-based Mendelian randomisation (SMR)

The SMR and HEterogeneity In Dependent Instrument (HEIDI) tests (Zhu *et al.*, 2016) are implemented in the SMR software (see URLs). We applied the SMR method to summary-level GWAS data and the sex-stratified X-chromosome eQTL data generated from in our analyses (UKB and CAGE, respectively) to test for pleiotropic associations between 1,639 X-linked gene expression probes and 20 complex trait phenotypes in SMR analysis. A total of 135, 113 and 66 probes with at least one *cis*-eQTL at GWS threshold P<5.0×10^−8^ were retained in male and female and in a combined *cis*-SMR analysis, respectively. SMR analysis in *trans* regions was performed with combined data and 74 probes with *trans*-eQTLs P_eQTL_<5.0×10^−8^ were included. A reference for LD estimation was a random sample of 100,000 unrelated UKB females of European ancestry. Trait-gene associations were identified using a significance level of P_SMR_<3.0×10^−5^ (i.e 0.05/1,639) for SMR analysis. These associations were then tested for evidence of linkage, rather than pleiotropy/causality, using the HEIDI test, which tests for heterogeneity in the effect estimates of the exposure on the outcome at SNPs in LD with the top associated eSNP under the null hypothesis of no heterogeneity. Gene-trait associations with P_HEIDI_>0.05 were selected.

#### Estimating the effect size ratio and dosage compensation coefficient (DCC)

We refer to the effect size ratio as the ratio of M/F per-allele effect estimates for a single trait-SNP association. The corresponding standard errors are estimated as,

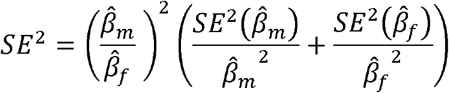

As before, 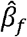 and 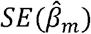 are the M/F per-allele effect estimates, and 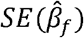 and 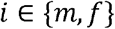 are the corresponding standard errors, respectively. To compare the per-allele effect estimates across all conditionally independent trait-associated SNPs (complex trait analysis) and top eQTLs (gene expression analysis) identified in the discovery datasets, we calculated an effect size regression coefficient (DCC) by regressing the per-allele effect estimates in males onto females weighted by inverse of the variance of male-specific estimates, and extracting the slope estimate and corresponding standard error. The estimates from sex-stratified XWAS, rather than joint effect estimates from the GCTA-COJO analysis were used for estimating DCC in the UKB traits. DCC is expected to take on values between 1 and 2, where DCC of 1 indicates that, on average, the effect sizes in males and females are equal (i.e. no DC or escape from XCI), and DCC of 2 indicates that, on average, the effect sizes in males are twice that of females (i.e. full DC).

#### X-chromosome gene inactivation status

To determine X-chromosome inactivation status, we downloaded annotation from the “Reported XCI status” column in Supplementary Table 13 of (Tukiainen, A.-C. Villani, *et al.*, 2017) and mapped gene expression probes to XCI status using the gene name. A total of 683 X-linked transcripts were available, where transcripts were classified as either “Escape” (82 transcripts), “Variable” (89 transcripts), “Inactive” (392 transcripts) or “Unknown” (120 transcripts). For each SNP in UKB dataset we determine if it is physically located within a gene to infer the presumable gene and its inactivation status for independent GWS SNPs.

## Full detail of Methods and Materials can be found in the Supplementary Methods and Material

### Supplementary Methods and Materials

#### Theoretical framework

Following (Lee *et al.*, 2018), the genetic variance contributed by an X-chromosome SNP, under the assumption of Hardy-Weinberg equilibrium (HWE), in females is,

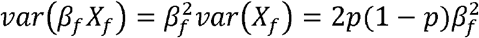

where *β*_*f*_ is the per-allele effect estimate from a regression of SNP, *X*_*f*_, on phenotype, *Y*_*f*_, with *X*_*f*_ ∊ {0,1,2}; and *p*, the minor allele frequency. Similarly, in males,

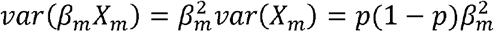

where, *β*_*m*_ is the per-allele effect estimate from a regression of SNP, *X*_*m*_ on phenotype, *Y*_*m*_, with *X*_*m*_ ∈ {0,1}. Dosage compensation can be parameterised as *β*_*m*_ = *dβ*_*f*_ where 1 ≤ *d* ≤ 2 in general,

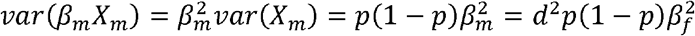

Under a full dosage compensation model (*d* = 2), *β*_*m*_ = 2*β*_*f*_ and,

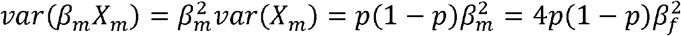

That is, the variance contributed by a X-linked SNP in males is twice that of females. Under a no dosage compensation model (*d* = 1), *β*_*m*_ = *β*_*f*_ and,

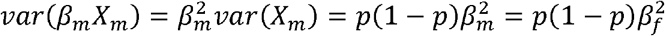

That is, the variance contributed by a X-linked SNP in males is half that of females. Further, we can estimate *d* (i.e. dosage compensation ratio) by exploiting the following relationship,

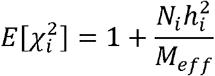

for *i* ∊ {*m*, *f*}, where, 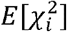 is the expected mean 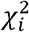 statistic for a gene; *N*_*i*_ is the sample size; 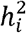 is the proportion of variance explained by X-chromosome SNPs; and *M*_*eff*_ is the effective number of X-chromosome SNPs. Rearranging for 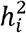 and taking the ratio 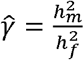, we get,

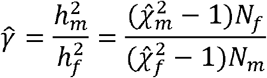

where 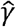 ranges between 0.5 (i.e. no dosage compensation) and 2 (i.e. full dosage compensation). Finally, the expectation of the cross-product of the *z*-statistics from the male and female analyses, 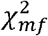 is,

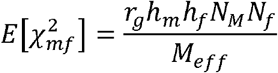

where *r*_*g*_ is the genetic correlation between males and females. Rearranging,

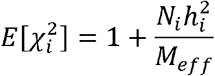

for 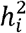 and substituting, we get,

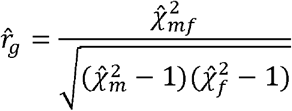

### UK Biobank Data

#### Sample selection

The complex trait analysis was conducted utilizing the UK Biobank (UKB) data (available to researchers upon application; see URLs). We inferred ancestries of 488,377 genotyped participants of the UKB as described in (Yengo *et al.*, 2018), and a dataset of European-ancestry individuals that met our sample quality inclusion criteria (N=455,605) was taken forward for the analysis. The samples were excluded according to UKB provided information if: (i) the genetically inferred gender was inconsistent with the submitted gender, (ii) there was evidence for putative sex chromosome aneuploidy, (iii) samples were reported as heterozygosity and missingness outliers, (iv) were excluded from kinship inference, or if participants have withdrawn their consent for using the data.

#### Genotype data

The imputed genotypes for both autosomes and X-chromosome pseudo-autosomal (PAR, coded as chromosome 25) and non-PAR (coded as chromosome 23) regions are available as a part of the UKB Version 3 release of the genotype data. Individuals were genotyped on either Affymetrix UK BiLEVE Axiom (N=50,000) or the Affymetrix UK Biobank Axiom^®^ array (N=450,000). The genotypes were imputed to UK10K+1000GP3 and HRC reference panels and include both SNPs and small indels (Bycroft *et al.*, 2017). We further hard-called the provided genotype probabilities (chromosomes 1-22, 23 and 25) of non-multiallelic markers with info-score > 0.3, treating the calls with uncertainty > 0.1 as missing, and keeping the markers which meet our quality control criteria in the set of unrelated European individuals (HWE test P<10^−6^ and missing call rate <5%). The heterozygous calls in non-PAR region of the X chromosome male genotypes were set to missing. To avoid deflation of heritability estimates on the X chromosome we only analyse the markers with MAF>0.01 in our full sample of European participants. We estimate allele frequencies (AF) of the X-chromosome markers for both sexes and keep the common set of 6,871 PAR and 253,842 non-PAR SNPs.

#### Phenotype selection

A total of 20 complex traits were selected for the analysis in the UKB. All analyses as well as phenotype adjustment were performed on a sex-specific basis. The phenotypes were adjusted for covariates and the residuals were transformed to sex-specific z-scores (mean=0, variance=1) with the phenotype measure values over 6 standard deviations (SD) away from the mean previously removed from the analysis. For individuals with repeated measures of the phenotype, we estimated the mean value of the observed measures after outlier removal procedure for each assessment visit and used mean age across the visits as a covariate. For each trait the UK Biobank variable identifiers, available sample sizes and covariates are presented in **Supplementary Table 1a,** as well as the minimum, maximum and mean values of the raw phenotype measures and the standard deviations of the phenotype after adjustment for trait-specific covariates. The discrete phenotypes (educational attainment, smoking status, skin and hair colours) were treated as quantitative (**see Supplementary Table 1b for description of the categories**) in our association analysis.

### Consortium for the Architecture of Gene Expression (CAGE) data

#### Gene expression and X-chromosome genotype data

Gene expression and X-chromosome genotype data were available in a subset of N=2,130 individuals (N=1,084 males, N=1,046 females) from the Consortium for the Architecture of Gene Expression (CAGE), a study examining the genetic architecture of gene expression in a mixture of pedigree and unrelated individuals (Lloyd-jones *et al.*, 2017). This subset of individuals comes from three cohorts with genotype data on the X chromosome (Powell *et al.*, 2012, 2013; Kim *et al.*, 2014; Leitsalu *et al.*, 2015), and are of European ancestry, as identified by principal component analysis with the HapMap3 populations. Further details are provided in (Lloyd-jones *et al.*, 2017).

#### Quality control of gene expression data

RNA was collected from whole-blood samples in each cohort and gene expression levels quantified using the Illumina Whole-Genome Expression BeadChips (HT12 v.3 and HT12 v.4). A total of 38,624 gene expression probes were common to all cohorts. Gene expression quality control and normalisation was performed in each cohort separately before concatenation. This included variance stabilisation and quantile normalisation to standardise the distribution of expression levels across samples. To remove hidden and known experimental confounders, gene expression levels were then adjusted for a mean of 39/50 PEER factors (Stegle *et al.*, 2010, 2012) across the three cohorts that were not associated with sex (P_sex_>0.05) in order to preserve the effect of sex on expression and where available, measured covariates such as age, cell counts, and batch effects. Residuals for each cohort were then standardised to z-scores and concatenated across cohorts. The concatenated gene expression dataset was further adjusted for 18/50 PEER factors that were not associated with sex (P_sex_>0.05) and standardised to z-scores. A total of 36,267 autosomal and 1,639 X-chromosome gene expression probes (corresponding to 26,384 and 1,138 unique genes, respectively) that unambiguously mapped to the genome formed our final gene expression dataset. This included a total of 28 PAR X-chromosome gene expression probes.

#### Quality control and imputation of genotype data

Genotype data was acquired using different genotyping platforms for each cohort, with quality control performed within each cohort before concatenation. Details for autosomal quality control and imputation are provided in (Lloyd-jones *et al.*, 2017). Briefly, autosomal SNPs were imputed to the 1000 Genomes Phase 1 Version 3 reference panel (Altshuler *et al.*, 2012) within each cohort and concatenated resulting in 7,763,174 SNPs passing quality control, which included filtering SNPs for minor allele frequency (MAF) <0.01, HWE test P<10^−6^, and imputation info score <0.3. This set of imputed autosomal SNPs was further filtered to 1,066,905 HapMap3 SNPs that were common to all three cohorts. This set of imputed autosomal SNPs formed our final dataset. For each cohort, we used the Sanger Imputation Server (https://imputation.sanger.ac.uk/) to impute SNPs on the non-PAR of the X chromosome to the Haplotype Reference Consortium (HRC, release 1.1) (McCarthy *et al.*, 2016), using the EAGLE2+PBWT pre-phasing and imputation pipeline (Durbin, 2014; Loh *et al.*, 2016). Pre-imputation checks included ensuring all alleles are on the forward strand, and coordinates and reference alleles are on the GRCh37 assembly. Pre-imputation quality control included filtering X-chromosome genotyped SNPs for MAF<0.01, HWE test P<10^−6^ within females, SNP missingness call rate >2%, and genotyped SNPs that are not in the HRC reference panel. A total of 1,228,034 X-chromosome SNPs were available following imputation in each cohort. Post-imputation quality control within cohort included filtering imputed X-chromosome SNPs for MAF<0.01, HWE test P<10^−6^ within females, imputation info score <0.3, and multiallelic SNPs. A total of 306,589 imputed X-chromosome SNPs were common to all cohorts and formed the concatenated dataset. We performed further quality control of the concatenated dataset by filtering imputed X-chromosome SNPs for missingness call rate >2%. A total of 190,506 imputed X-chromosome SNPs remained. Additional post-imputation quality control on the concatenated dataset included a comparison of allele frequencies between males and females, which led to the exclusion of 261 SNPs with MAF differences of >0.05 between sexes. A total of 190,245 imputed X-chromosome SNPs formed our final dataset.

### Genotype Tissue Expression (GTEx) data

We used the Genotype Tissue Expression project (GTEx v6p release) dataset comprised of RNA-seq data from 39 non-diseased tissue-types for which a sex covariate was available in N=449 deceased human donors as an external validation of our X-chromosome *cis*-eQTL results across multiple tissue-types. The fully-processed, normalised and filtered RNA-seq GTEx v6p data were downloaded from the GTEx Portal (https://www.gtexportal.org/home/datasets) along with corresponding covariate files. X-chromosome imputed SNP data was obtained from dbGap (Accession phs000424.v6.p1). Briefly, gene expression normalisation included filtering for transcripts with at least 10 samples with RPKM >0.1 and raw read counts greater than 6, quantile normalisation within tissue, and inverse quantile normalisation for each transcript. Sample outliers were identified and excluded using a correlation-based statistic described in (Wright *et al.*, 2014), and samples with less than 10 million mapped reads were excluded. Further details can be found in (Consortium, 2017). Quality control of the X-chromosome imputed SNP data included filtering for MAF<0.05, HWE test P<10^−6^ within females, imputation info score <0.4, and multiallelic SNPs. A total of 127,808 imputed SNPs in the non-PAR of the X chromosome were included in our analysis. We restricted our analyses to 22 tissue samples for which within tissue sample size was greater than N=50 in both males and females (**Supplementary Table 10**). Sample sizes per tissue ranged from N=124 in colon (sigmoid) to N=361 in muscle (skeletal) with a mean of N=226 across the 22 tissues. The proportion of males and females within each tissue ranged from 34% females in heart (atrial appendage) to 44% females in adrenal gland, with a mean of 38% females across all 22 tissues. A total of 1,121 X-linked transcripts (including 31 PAR transcripts) were expressed in at least one tissue of the 22 tissues. The number of X-linked transcripts identified as expressed in each tissue ranged from 726 in pancreas to 916 in thyroid, with a mean of 808 across all 22 tissues (**Supplementary Table 10.**).

### Statistical Analysis

#### GWAS

To determine the DC ratios across 20 complex traits and to compare effect sizes of genome-wide significant X-chromosome markers on those phenotypes, we analyse the results of X-chromosome wide analysis (XWAS) (both PAR and non-PAR) performed on a sex-specific basis using BOLT-LMM v2.3 (Loh *et al.*, 2018) in the full set of UKB European males (N_m_=208,419) and females (N_f_ =247,186). We include a set of HapMap3 SNPs (MAF>0.01 and pairwise R^2^<0.9 in the window of 1000 SNPs) in the mixed model to correct for the population stratification and to account for relatedness. This set of model SNPs (M=561,572) includes autosomal markers, 12,508 non-PAR and 205 PAR SNPs on the X chromosome. All other X-chromosome SNPs are fixed effects and tested for association using linear regression.

#### Combined analyses

The choice of the optimum meta- and combined analyses depends on the assumptions of dosage compensation and the genotype coding in males (see Supplementary information in Lee *et al.*, 2018). While the true extent of dosage compensation is not known, its effect can be parameterised as *β*_*m*_ = *dβ*_*f*_, with *d* being a dosage compensation parameter (*d* = 1 for no dosage and *d* = 2 for full dosage compensation). In the sex-stratified analysis we regress a phenotype on a genotype variable, where *X*_*f*_ {0,1,2} for females and *X*_*m*_ɛ{0,c}in males, with *c* = 1 in the no DC analysis or *c* = 2 in the full DC analysis (i.e. assuming full random X-inactivation). When *c* = 1, we estimate per-allele effects in males. From the Eq. 4.6 and 4.7 in (Lee *et al.*, 2018), it follows that an optimum meta-analysis of the estimates from the sex-stratified analysis is only unbiased when *d* = *c*. That is, under a no DC model, the meta- and combined analyses will be unbiased when using per-allele effect estimates in males (*c* = 1), while under a full DC model, they are unbiased when the effect estimates in males are from an association analysis where the male genotypes coded as diploid (*c* = 2). Since the results from our sex-stratified analysis are largely consistent with expectations from full dosage compensation, we perform an inverse variance weighted meta-analysis for complex traits using the male effect size estimates from the diploid analysis to obtain the joint estimates of the SNP effects, and in the combined analyses of gene expression traits we code males as diploids.

#### Sexual dimorphism in gene expression

Sexual dimorphism in gene expression was examined with a mixed linear regression model implemented in the GCTA software package (Yang, Lee, *et al.*, 2011). Here, we tested for sex differences in gene expression for 1,639 X-linked gene expression probes. Gene expression was modelled as,

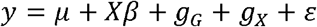

where *y* is a N × 1 vector of gene expression intensity levels; is the mean expression levels; *β* is the regression coefficient for the fixed sex covariate, *X*, with males coded as 1 and females coded as 2; *g*_*G*_ is an N × 1 vector of the total genetic effects of the individuals with 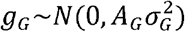, where *A*_*G*_ is interpreted as the autosomal GRM between individuals calculated from 1,066,905 HapMap3 SNPs; *g*_*X*_ is an N × 1 vector of X-linked genetic effects with 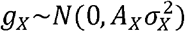 where *A*_*X*_ is a GRM calculated from 190,506 imputed X-chromosome SNPs; and 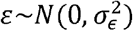 is the residual. We used the Wald statistic to assess significance, and calculated a P-value by comparing the test statistic to a *χ*^2^-distribution with one degree of freedom.

#### X-chromosome *cis*-eQTL analysis

To investigate the X-chromosome genetic control of gene expression, we modelled gene expression levels as a linear function of the number of reference alleles in a linear mixed regression model, in males and females separately and in a combined analysis, using the GCTA software package (Yang, Lee, *et al.*, 2011). The model for each gene expression probe can be written as,

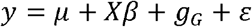

where, *y* is a N × 1 vector of gene expression intensity levels, with sample size N; *β* is a vector of fixed effect estimates for the indicator variable for the genotype, *X*; *g*_*G*_ is an N × 1 vector of the total genetic effects of the individuals with 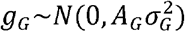, where *A*_*G*_ is interpreted as the autosomal genetic relationship matrix (GRM) between individuals calculated from the 1,066,905 HapMap3 SNPs; and 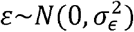 is the residual. Since our interest is in testing for the association between X-chromosome SNPs and gene expression, this is equivalent to a leave-one-chromosome-out analysis (J. Yang *et al.*, 2014). To assess significance, we calculated a likelihood ratio test statistic and calculated a P-value by comparing the test statistic to a *χ*^2^-distribution with one degree of freedom. We accounted for multiple testing for both the number of X-chromosome SNPs and the number of gene expression probes tested using the Bonferroni method. For each gene expression probe, eQTLs were defined as the top associated X-chromosome SNP that satisfies the Bonferroni significance threshold of P<1.6×10^−10^ (i.e. 0.05/(1,639×190,245) in the discovery sex. The XCI status (escape/variable or inactive) for the identified eQTLs were assigned by mapping gene expression probes to XCI status using the gene name from (Tukiainen, A.-C. Villani, *et al.*, 2017).

#### Autosomal *cis*-eQTL analysis

We compared results from our sex stratified X-chromosome *cis*-eQTL analysis to the autosome by performing an autosomal *cis*-eQTL analysis in males and females, separately. Here, we model autosomal gene expression levels as a linear function of the number of reference alleles for autosomal SNPs on the same chromosome using the GCTA software package (Yang, Lee, *et al.*, 2011). Each autosomal gene expression probe is modelled in the same way as described above. We identified eQTLs as probe-SNP pairs with P<10^−10^ in the discovery sex.

#### X-chromosome *cis-*eQTL analysis in GTEx

We modelled gene expression as a linear function of the number of reference alleles in a linear regression model for males and females separately using PLINK (Purcell *et al.*, 2007). The model for each X-chromosome transcript can be written as,

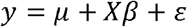

where, *y* is a N × 1 vector of gene expression intensity levels, with sample size N; *β* is a vector of fixed effect estimates for the for the indicator variable for the genotype, *X*; and 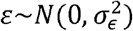 is the residual. The model was adjusted for three genotyping principal components (PCs) and PEER factors, which captures batch effects and latent experimental confounders in the gene expression data. Following (Consortium, 2017), a total of 15 PEER factors were included in the model for total sample sizes N<150, 30 PEER factors for total sample sizes 150≤N<250, and 35 PEER factors for total sample sizes N≥250. To assess significance, we calculated a t-statistic and calculated a P-value by comparing the test statistic to the t-distribution. We identified eQTLs as transcript-SNP pairs that satisfied the within tissue Bonferroni significance threshold, which accounts for both the number of X-linked transcripts and X-chromosome SNPs tested in each tissue in the discovery sex (**see Supplementary Table 10**). DCC was estimated in each of the 22 tissue-types as previously described. The XCI status (escape/variable or inactive) for the identified eQTLs in each tissue was assigned by mapping transcript gene identifiers from (Tukiainen, A.-C. Villani, *et al.*, 2017). We tested for enrichment of escape/variable status in each tissue using a hypergeometric test. As the proportion of males and females within each tissue is highly skewed towards males, sensitivity analysis included randomly removing male samples from the analysis so that the proportions match that of females within each of the tissues. This is repeated 100 times, with DCC calculated across the 100 replicates. Finally, we identified the top eQTLs among all tissues in the discovery sex, and extracted the corresponding eQTL from the same tissue in the other sex. DCC is calculated as previously described.

